# Morphological and physiological effects of a single amino acid substitution in the patatin-like phospholipase CapV in *Escherichia coli*

**DOI:** 10.1101/2020.11.22.387274

**Authors:** Fengyang Li, Heike Bähre, Manfred Rohde, Ute Römling

## Abstract

In rod-shaped bacteria morphological plasticity occurs in response to stress, which blocks cell division to promote filamentation. We demonstrate here that overexpression of the patatin-like phospholipase variant CapV_Q329R_ but not CapV causes pronounced *sulA*-independent pyridoxine-inhibited cell filamentation and restriction of swimming and flagella production of *Escherichia coli* K-12 derivative MG1655. Mutational analyses of CapV_Q329R_ indicated conserved amino acids in canonical patatin-like phospholipase A motifs, but not the nucleophilic serine to be required for the observed phenotypes. Furthermore, CapV_Q329R_ alters rdar biofilm formation including expression of the biofilm activator CsgD. Moreover, commensal and pathogenic *E. coli* strains and *Salmonella typhimurium* also responded with cell filamentation and alteration in biofilm formation. In conclusion, this work identifies the CapV variant CapV_Q329R_ as a pleiotropic regulator, emphasizes a scaffold function for patatin-like phospholipases and highlights the role of a single amino acid change for the evolution of protein functionality.

## Introduction

Bacteria are traditionally defined by their cell shape such as rod, coccus, spiral or filamentous and exist as unicellular or multinucleated cells (*1*). Shaped differently, many bacterial species display extensive morphological plasticity in response to environmental cues such as severe stress (*2, 3*). This morphological variation is often reversible suggesting an altered physiological cell status or epigenetic modulation of the genetic information upon signal perception or manifested threat. For example, upon escape from intracellular biofilms during late infection of bladder cells, rod-shaped uropathogenic *Escherichia coli* (UPEC) can reversibly transform from a rod with a length of 2 - 4 μm into a filament of up to 70 μm in length (*4*). In nature, *Caulobacter crescentus,* a freshwater curved rod shaped bacterium, can develop 20 μm long helical filaments which are suggested to reach out of biofilms for nutrient acquisition (*5, 6*). Other environmental bacteria such as *Epulopiscium fishelsoni* and *Thiomargarita namibiensis* develop up to 750 μm long filaments (*7*).

Upon nutrient deficiency (*6, 8*), antibiotic treatment (*9–11*) DNA damage (*12–14*) and other various stress conditions, bacterial cell filamentation is a consequence of activation of the SOS response system. Thereby, mainly functionality of the essential cell division protein, the tubulin homolog FtsZ, which initiates the cell division process by forming a septal ring at the prospective invagination site, is impaired (*7, 15, 16*). Expression of *sulA,* encoding a major SOS inhibitor of septum forming FtsZ, eventually leads to cell filamentation (*16–19*). The subsequent arrest in growth enables the cells to eventually recover from otherwise deleterious damages before resuming growth.

However, there exist alternative *sulA*-independent stress-associated pathways that promote filamentation in *E. coli.* For example, upon treatment with cationic antimicrobial peptides, QueE, an enzyme involved in the queuosine tRNA modification pathway, blocks cell division and induces filamentation (*20*).

Development of filamentation can also serve alternative purposes. As a survival strategy, filamentation contributes to pathogenesis. Treatment with cell wall inhibiting β-lactam antibiotics in patients, can promote bacterial surface colonization by triggering *E. coli* filamentation (*21, 22*). In UPEC *E. coli,* filamentation can slow down phagocytosis by macrophages during infection and thus enhances bacterial survival upon this challenge by the host immune responses (*4, 17, 23*).

During cell division cell elongation, chromosome replication and segregation and cytokinesis (cell separation) are coordinated (*24–26*). In this complex process, dysregulation of critical components and regulators can promote cell filamentation (*25, 27–29*). For example, overexpression of the cell division inhibitor MinC, which prevents FtsZ septum formation at the cell poles by oscillation from pole to pole assisted by MinDE leads to filamentation (*30*). DamX, a membrane-spanning protein with a peptidoglycan binding SPOR domain, is required for reversible filamentation, colonization and pathogenesis of UPEC morphotype switching (*31*).

Motility is defined as the ability to actively move in liquid or on surfaces (*32*). Bacterial motility covers various modes of movement dependent on energetic requirements and the structural elements involved (*33*). As a common mode of motility propelling of flagellar filaments moves bacterial cells either in liquid medium by swimming motility or on a surface of semi-solid medium by swarming motility (*32, 34*). In *E. coli* and *S. typhimurium* bacterial motility, on the level of the class 1 transcriptional regulator FlhD4C2 and downstream, is tightly regulated by global regulatory signals such as c-di-GMP signaling and protein-protein interactions of enzymatically incompetent c-di-GMP related proteins (*34, 35*). Cyclic di-GMP is a ubiquitous bacterial intracellular second messenger that ubiquitously modulates the lifestyle change between motility and sessility (biofilm formation) (*36, 37*). While, in *E. coli* and *S. typhimurium*, flagella-based swimming and swarming motility are inhibited by c-di-GMP on the posttranslational level targeting flagella motor functionality (*34*), c-di-GMP activates rdar biofilm formation and expression of *csgD* coding for the major transcriptional regulator of biofilm formation. Besides the ubiquitous c-di-GMP signal, also other recently identified c-di-nucleotide second messengers modulate the sessility/motility life style transition. Such inhibits the second messenger cyclic AMP-GMP (3’3’-cGAMP) synthesized by the cyclic dinucleotide cyclase DncV (*38, 39*) motility and rdar biofilm formation in the commensal *E. coli* strain ECOR31 (*39*). *DncV* is part of a putative *capV-dncV-vc0180-vc0181* four-gene operon encoded by a right border accessory part of the horizontally transferred *Yersinia* high-pathogenicity island (HPI) (*40, 41*). Thereby, *capV* encodes a patatin-like phospholipase activated by 3’3’-cGAMP to induce growth retardation in *V. cholerae* (*42*). In this study, we show that upon overexpression, the variant CapV_Q329R_ causes substantial alterations in morphology on the single and multicellular level leading to extended *sulA*-independent cell filamentation and restricted swimming motility with premature flagella loss. Besides these morphological changes, which occur in commensal and pathogenic strains of *E. coli* and *S. typhimurium*, CapV_Q329R_ altered rdar biofilm formation and colony morphology. These findings demonstrate, as an example of rapid evolution of protein functionality, a single amino acid change in a patatin-like phosphodiesterase leads to the manipulation of various aspects of bacterial physiology independent of catalytic activity

## Results

### CapV_Q329R_ inhibits swimming motility of *E. coli* MG1655

The dinucleotide cyclase DncV synthesizes 3’ 3’-cGAMP to inhibit rdar biofilm formation and motility in the animal commensal strain *E. coli* ECOR31 (*39*). *DncV* is flanked by *V. cholerae* 7^th^ pandemic island-1 (VSP-1) homologs of *capV, vc0180* and *vc0181* (*38–40*) (fig. S1A). These genes constitute a putative 4.6 kbp four-gene operon which contributes to differential functionality of *dncV* in *E. coli* ECOR31 (*39*). To understand physiological functions of the four-gene operon (designed *78901*), we tested swimming motility of the *E. coli* K-12 derivative MG1655 upon overexpression of *78901* in semi-solid agar. Interestingly, overexpression of the four-gene operon *78901* from a plasmid significantly inhibited swimming motility at 37 °C and 28 °C (fig. S1B and C). Since the plate assay monitors both chemotaxis and motility, we assessed the production of cell associated extracellular flagellin upon overexpression of *78901.* While wild-type MG1655 showed pronounced flagellin production under our experimental conditions, which stems from polymerized flagella, overexpression of *78901* inhibited flagellin production (fig. S1D).

To this end, we could show that the variant CapV_Q329R_ caused suppression of swimming motility (as described in Supplemental results; Fig. 1A, B, fig. S2 and S3). To further characterize CapV_Q329R_ induced MG1655 swimming inhibition, we assessed the production of cell-associated flagella and flagellin upon overexpression of CapV_Q329R_. Visualization of bacterial cells by transmission electron microscopy (TEM) showed that overexpression of CapV_Q329R_ dramatically reduced the total number of flagella-producing cells as well as the number of flagella per cell after 6 h incubation at 37 °C (Fig. 1C and D). In agreement, visualization of flagella by Leifson stain showed cells with intact flagella after 4 h incubation at 37 °C, but almost no cell with flagella after 6 h upon CapV_Q329R_ overexpression (Fig. 1E). In congruence with the analysis by TEM and the Leifson stain, we observed inhibition of production of cell associated extracellular flagellin in a protein gel upon overexpression of CapV_Q329R_ (Fig. 1F and G). Cumulatively, these results indicate that CapV_Q329R_ suppresses swimming motility of MG1655 by (temporarily) inhibiting FliC production.

**Fig. 1.**
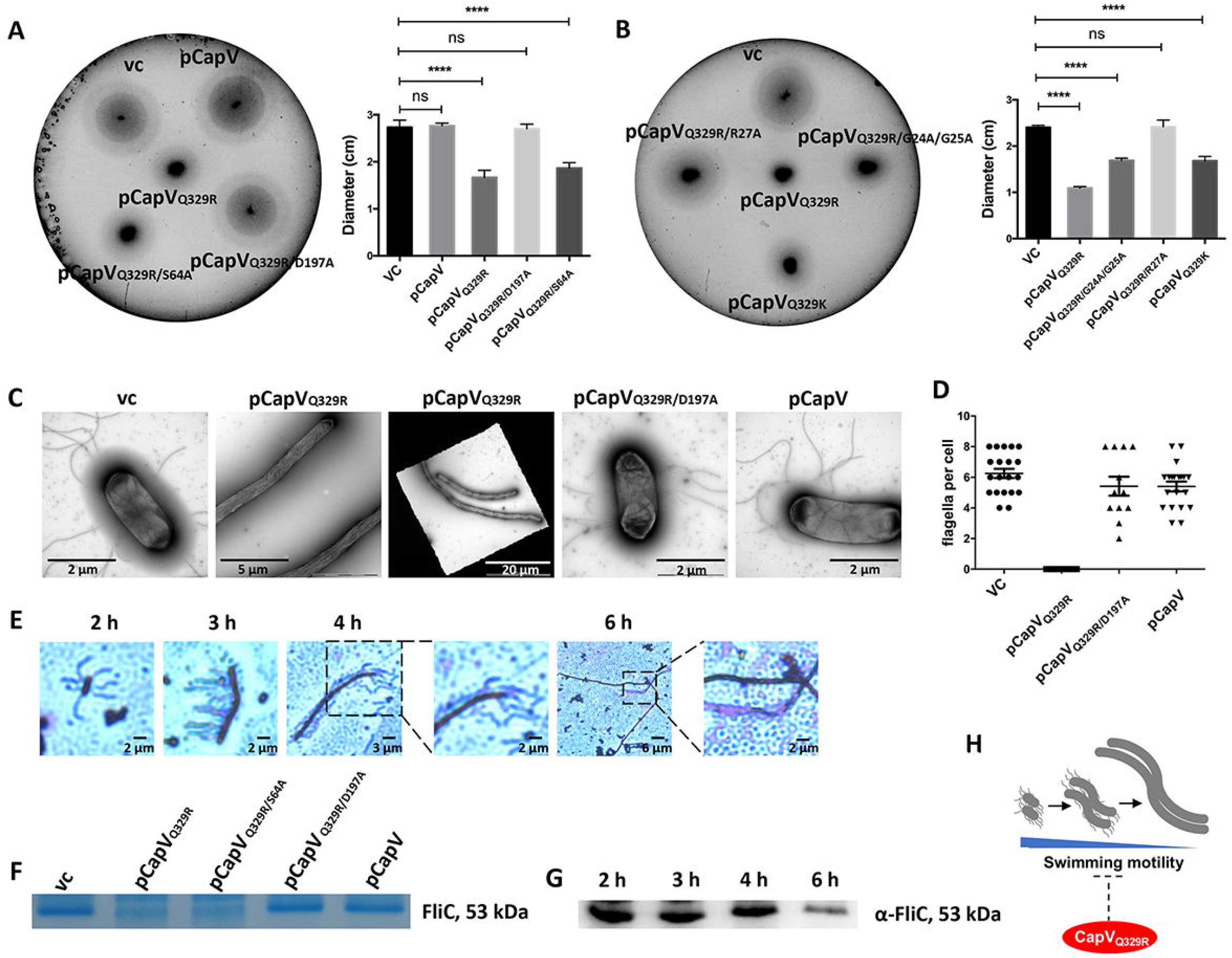
CapV_Q329R_ inhibits apparent swimming motility of *E. coli* MG1655. (**A** and **B**) Flagella-dependent swimming motility of wild type *E. coli* MG1655 and upon overexpression of CapV and its mutants. 3 μl of OD_600_ = 5 cells were inoculated into soft agar plates containing 1% tryptone, 0.5% NaCl and 0.25% agar and the swimming diameter was measured after 6 h at 37 °C. (**C**) Flagella production of a representative *E. coli* MG1655 cell and upon overexpression of CapV and CapV_Q329_R as observed by TEM. (**D**) Quantification of the number of flagella per cell upon overexpression of CapV and CapV_Q329R_ after visualization by TEM. Number of evaluated cells n=20. Cells were grown in TB medium for 6 h at 37 °C. (**E**) Production of surface-associated flagellin is down-regulated upon overexpression of CapV_Q329R_, but not upon overexpression of CapV wild type and mutants. Colloidal Coomassie staining of flagellin extracted from *E. coli* MG1655 vc, and expressing CapV_Q329R_. (**F** and **G**) Flagellin subunit FliC expression was decreased gradually upon expression of CapV_Q329R_. Cells were grown in TB medium at 37 °C, and samples were harvested at different time points in the growth phase for Western blot analysis of FliC and for flagella staining. (**H**) Proposed development of filamentation and flagella inhibition of *E. coli* MG1655 by CapV_Q329R_ over time. Bars represent mean values from three independent replicates with error bars to represent SD. Differences between mean values were assessed by two-tailed Student’s t-test (ns, not significant; *p < 0.05, **p < 0.01, and ***p < 0.001 compared to *E. coli* MG1655 VC). VC = pBAD28. pCapV = CapV cloned in pBAD28; pCapV_Q329R_ = CapV_Q329R_ cloned in pBAD28.

### CapV_Q329R_ promotes cell filamentation of *E. coli* MG1655

Furthermore, TEM analysis displayed the development of pronounced long thin cell filaments of MG1655 upon overexpression of CapV_Q329R_ when grown in tryptone broth (TB) medium (Fig. 1C). In contrast, the cell length of MG1655 cells overexpressing CapV wild type was only slightly elongated compared to the MG1655 vector control (VC) (Fig. 1C).

We wanted to assess the temporal development of cell filamentation throughout the growth phase upon induced expression of CapV_Q329R_ by light microscopy. Temporal follow-up indicated initial development of filamentation did not initiate before 2 h after commencement of CapV_Q329R_ expression. Subsequently, though, the cell length and the frequency of filamentation dramatically increased, whereby after 3 h almost all cells displayed as short filaments around 5 times the length of standard rod cells (Fig. 2A and B). After 6 h induction, CapV_Q329R_ expressing *E. coli* MG1655 cells were around 25 times longer than MG1655 VC cells; however, many filaments even were 50 times the length of a standard rod-shaped cell (Fig. 2B). Those long cells did not show any movement, while shorter filaments up to approximately 20 times the length still showed swimming motility (movie S1). At the opposite, overexpression of CapV wild type only slightly increased the cell length compared to MG1655 VC. Interestingly, after 22 h induction by 0.1% L-arabinose, short rod-shaped cells dominated again, which suggests filaments to restart cell division after L-arabinose depletion. In line with this hypothesis, induction of CapV_Q329R_ production by 0.2% L-arabinose did not cause reversion to rod-shape nor showed any movement after overnight culture (Fig. 2A, movie S2). On the other hand, upon transfer of filamentous cells to fresh TB medium without L-arabinose, emergence of rod-shaped cells was observed (Fig. 2C). The developmental process of filamentation with subsequent loss of flagella is displayed in Fig. 1H.

**Fig. 2.**
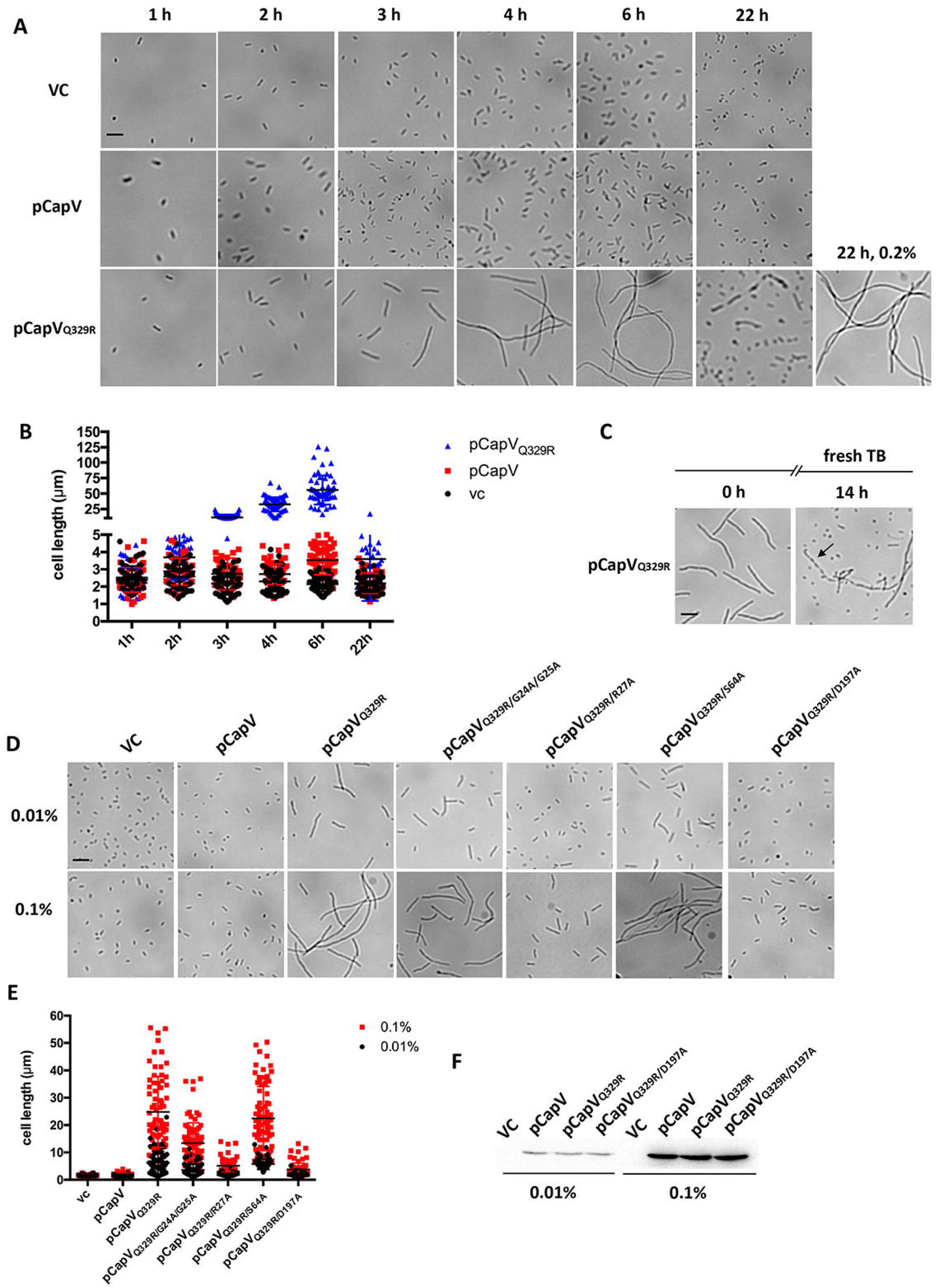
CapV_Q329R_ promotes cell filamentation of *E. coli* MG1655. (**A**) Light microscopy pictures of cell morphology of *E. coli* MG1655 and derivates in TB at different time points. (**B**) Quantification of cell length of MG1655 and derivates in TB at different time points. The quantification is based on results from at least three independent experiments with assessment of 70 cells from each group. (**C**) Recovery images of cells from a 3 h culture containing fresh TB medium. Arrowheads indicate invaginations at future division sites. (**D** - **F**) Light microscopy pictures (**D**), quantification of cell length (**E**) and protein expression level (**F**) of *E. coli* MG1655 and derivates with 0.01% and 0.1% L-arabinose induction in TB at 4 h. The quantification is based on results from at least three independent experiments with assessment of 70 cells from each group. Bar, 5 μm. VC = pBAD28.

Linearity of the filamentation phenotype of the MG1655 cells with CapV_Q329R_ production level was demonstrated by using a lower L-arabinose concentration. When incubated for 4 h with 0.01% L-arabinose, the low level CapV_Q329R_ expression created a mixed cell population with mild or no cell filamentation (Fig. 2D) in contrast to 0.1% L-arabinose where all cells became filamentous. Concomitantly, the CapV_Q329R_ expression level was higher with 0.1% L-arabinose than with 0.01% L-arabinose induction (Fig. 2D, E and F).

### CapV_Q329R_ induces asymmetrically positioned FtsZ rings and abnormal nucleoids in filaments

Upon cell division, FtsZ ring positioning at the nucleation site between nucleoids is coordinated with chromosome replication and nucleoid segregation (*28, 43, 44*). Thereby, cell division arrest and filamentation can be induced by DNA damage and nucleoid occlusion (*13, 24*). We analyzed FtsZ ring positioning in an *E. coli* K-12 MG1655 derivative with FtsZ-GFP fusion protein expression from the chromosome (BS001, (*45*)) upon CapV and CapV_Q329R_ overexpression compared to wild type. Furthermore, we assessed nucleoid positioning in these cells by staining the cells with DAPI and immediately subjected them to fluorescence microscopy on agarose pads to visualize DNA, septa, and/or the cell envelope. In the FtsZ-GFP derivative overexpressing wild type CapV, we observed clearly visible septa with a correctly positioned FtsZ ring and a single nucleoid in non-dividing cells and two fully replicated and/or segregated nucleoids in dividing cells (Fig. 3A).

**Fig. 3.**
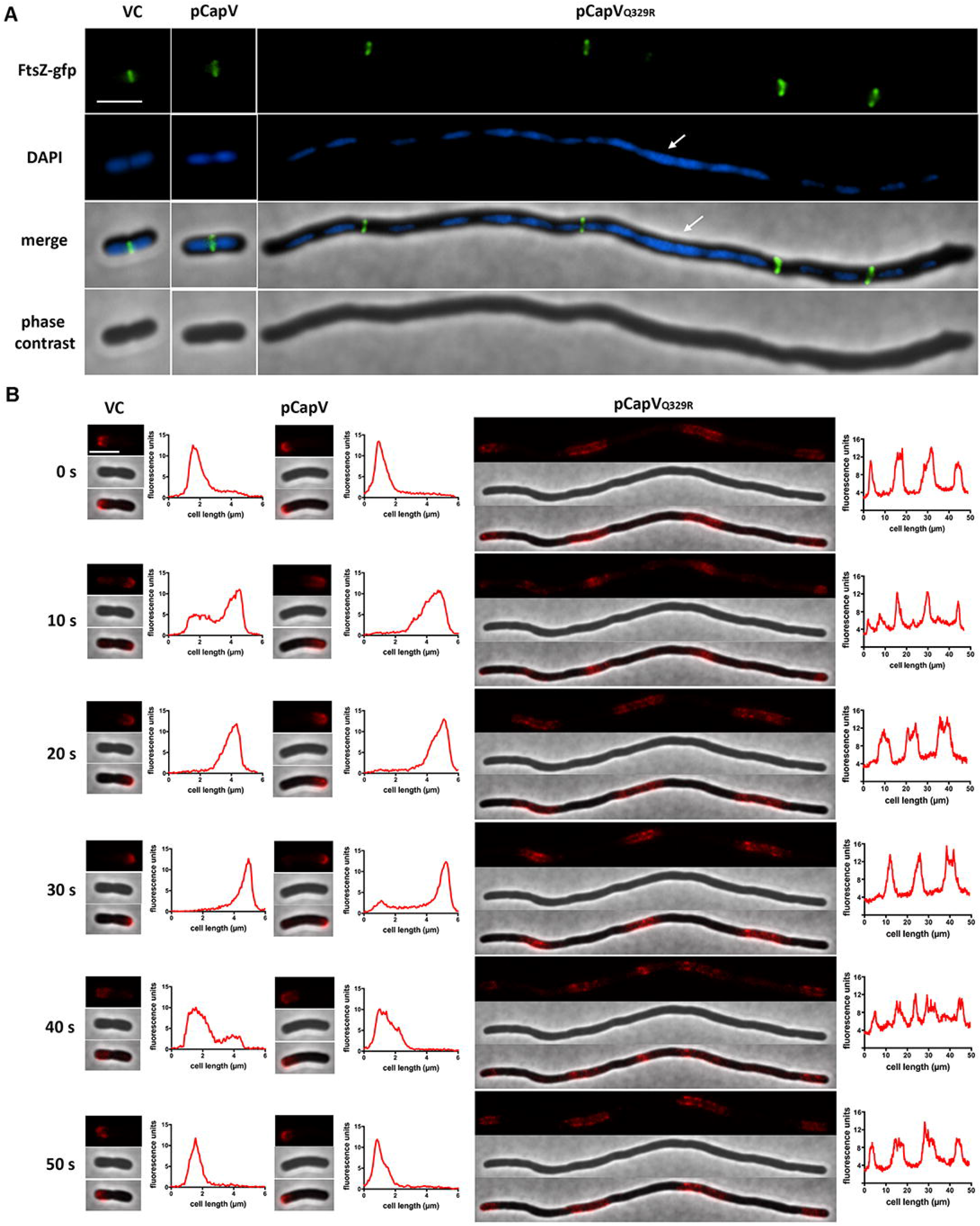
Chromosomal segregation, but not FtsZ ring formation, is impaired in filamentous cells. (**A**) Phase contrast and fluorescence images of FtsZ-GFP expressing cells (BS001). Cells were cultured in TB medium at 37 °C for 4 h, stained with DAPI and assessed immediately under fluorescence microscopy. Large fragments of unsegregated nucleoids are indicated by white arrows. Bar, 3 μm. (**B**) Time-lapse analysis of mCherry-MinC expressing cells (PB318). A representative elongating cell is displayed. Graphs on the right of fluorescence images display the line profiles of fluorescent signals emanating from the cell. Arbitrary fluorescent units are obtained, analyzed by the Fiji ImageJ 1.8.0 software (70) and are plotted on the y axis; cell length (in μm) is plotted on the x axis. Bar, 3 μm.

In contrast, upon CapV_Q329R_-induced cell elongation, most of the filaments displayed smooth contours with no visible septa (Fig. 3A, fig. S4A), suggesting that the block in cell division is due to inhibition of septation. However, one or two septa were occasionally observed in a few filaments (fig. S4A and B). Unexpectedly, we found that most of the CapV_Q329R_-induced filamentous cells contain abnormally shaped or positioned nucleoids. The abnormal nucleoid-staining patterns included nucleoids that are asymmetrically positioned in the filament, extended nucleoids or decondensed nucleoids occupying an extensive part of the filament (Fig. 3A, fig. S4B), suggesting the filamentous cells induced by CapV_Q329R_ overexpression are (partially) defective in chromosome segregation and nucleoid condensation. However, a few filaments contained normal nucleoids, evenly positioned throughout the filament.

In contrast, distinct FtsZ rings were observed in most of the filaments, but the distance between two FtsZ rings differed leading to an unequal number of rings in filaments of similar length. After 4 h induction of CapV_Q329R_, the majority of the filamentous cells contain 3 rings, with an average cell length of 31.2 μm (n = 102) with multiple segregated and/or unsegregated nucleoids distributed between the Z-rings. Interestingly, both the distance between two adjacent Z-rings and the number of partitioned nucleoids varies dramatically among the filamentous cells (Fig. 3A).

Abnormal FtsZ expression levels induces cell filamentation (*46*). Immunoblot analysis showed slightly lower FtsZ protein levels between vector control and CapV_Q329R_ overexpression cells (fig. S4C). Whether this (partial) decrease in FtsZ levels is responsible for CapV_Q329R_-induced filamentation needs to be further investigated.

### CapV_Q329R_-induced cell filamentation is *sulA*-independent

Previous studies showed that SulA blocks cell division by direct interaction with cell division core component FtsZ during the bacterial SOS response (*7, 29*). To investigate whether CapV_Q329R_ induced cell filamentation of MG1655 is caused by the activation of *sulA,* we analyzed cell morphology upon CapV_Q329R_ overexpression in a *sulA* mutant (*47, 48*). We found that CapV_Q329R_ induced the same filamentous cell phenotype in the *sulA* mutant as in MG1655 wild type (fig. S4D), suggesting that CapV_Q329R_-induced cell filamentation is *sulA*-independant.

### MinC oscillation takes place in filaments

Many factors are required for the regulation of cell division. Among them is MinC, which oscillates from pole to pole in order to prevent FtsZ ring formation. Upon overexpression MinC causes cell filamentation. We investigated the mobility behavior of MinC using time-lapse fluorescence microscopy (Fig. 3B). In rod-shaped wild type cells, we observed that MinC oscillates between cell poles, moving a large fraction of the total fluorescence signals along the cell as described previously (*49*). A complete oscillation cycle lasted about 50 s under our experimental conditions. Furthermore, we found that the oscillation of the FtsZ-ring inhibitor MinC, is not affected in CapV_Q329R_-induced filaments (Fig. 3B) (PB318, (*49*)). MinC displayed various fluorescence fraction peaks ranging from 1 up to 5 depending on the length of the filaments (Fig. 3B, fig. S4E). Moreover, the fluorescence peaks were present within and at both or none of the poles in most of the filaments. Within one single filamentous cell, the number of the apparent fluorescence fractions followed a three-step principle: n n - 1 n (n ≥ 2, located at both poles) or n n +1 n (n ≥ 1, located within cell), during an oscillation periodicity of about 50 s, similar to that observed for the rod-shaped vector control and CapV wild type overexpression cells (Fig. 3B, fig. S4E). According to our knowledge, this is the first report on MinC oscillation in filamentous *E. coli* cells.

### Overexpression of CapV_Q329R_ reduces cell numbers

CapV has been identified as a patatin-like phospholipase which causes growth retardation upon binding to 3’3’-cGAMP in *V. cholerae* El Tor (*42*). In *E. coli* MG1655 though, overexpression of CapV_ECOR31 wild type did not affect cell division during the entire growth phase. Though, no effect was observed during the first 3 h of growth in liquid culture, overexpression of CapV_Q329R_ induced a mild growth arrest after 6 h (Fig. 4A). As extensive filamentation might not permit a direct correlation between OD and cell number, we tested cell viability by spotting MG1655 cells on agar plates. Compared to MG1655 vector control and cells expressing CapV approximately 50% of the CFU were obtained for MG1655 cells expressing CapV_Q329R_ (Fig. 4B). Consistently, cells stained with SYTO 9 for viability and propidium iodide (PI) for cell dead showed that CapV_Q329R_ production induced an approximately 50% decrease in cell viability of *E. coli* MG1655 after 6 h, while it did not cause cell death after 3 h (Fig. 4C, fig. S5A and B). Of note, one filament could contain seemingly live and dead cells. Interestingly, the uptake of SYTO 9 into individual cells varied widely in particular in CapV expressing cells, which showed no decrease in cell viability.

**Fig. 4.**
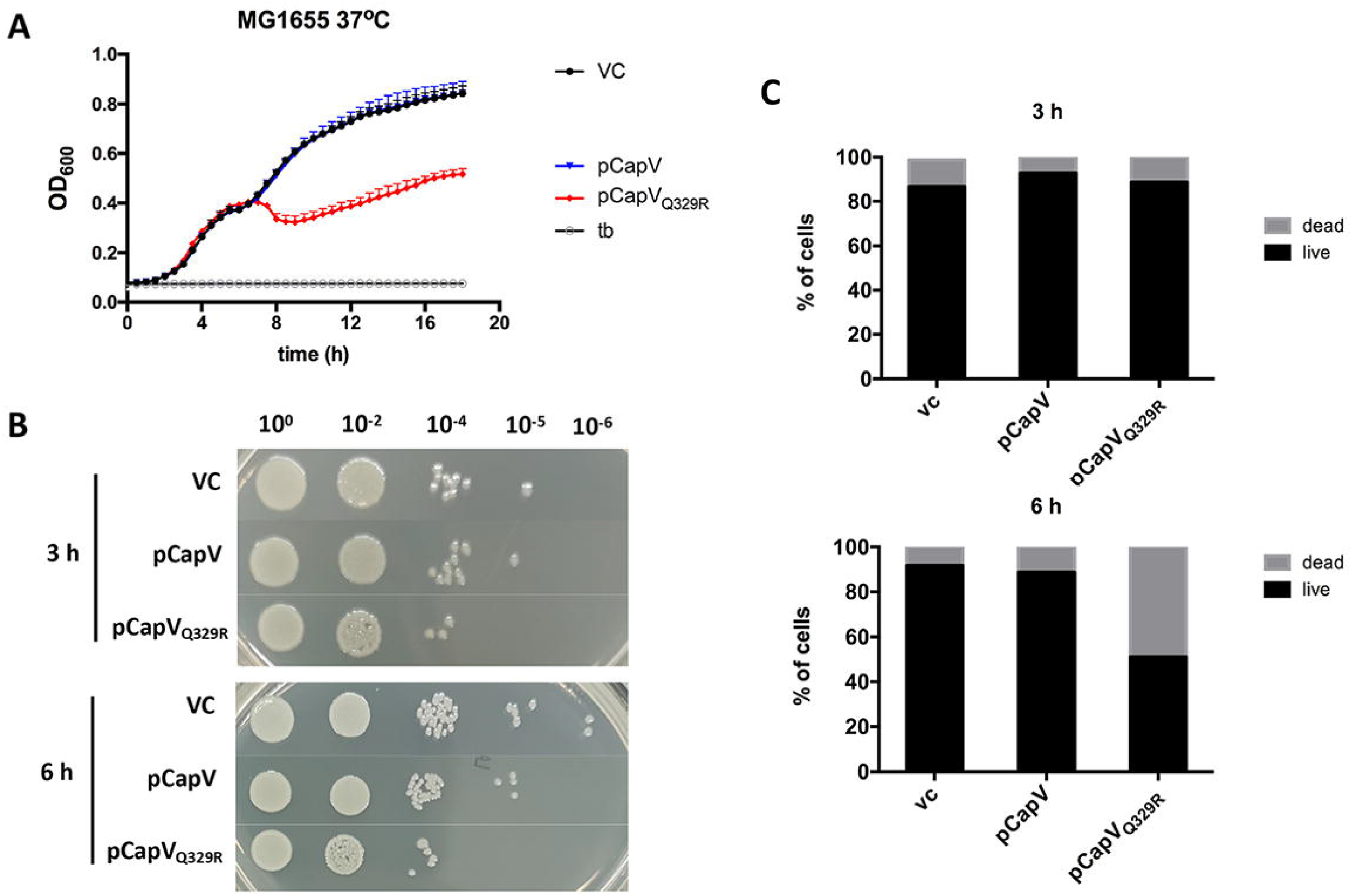
CapV_Q329R_ is cytotoxic to *E. coli* MG1655 in stationary phase. (**A**) Growth curves of MG1655 upon CapV and CapV_Q329R_ overexpression induced by 0.1% L-arabinose at 37 °C. Each data point represents the mean ± SD of six biological replicates. (**B**) Colony spotting assay on agar plates. Cells were grown at 37 °C and harvested at different time points. Cell viability determined by spotting serial dilutions (10^0^ – 10^-6^) on LB plates to assess colony forming units. (**C**) Quantification of Live/Dead staining of *E. coli* MG1655 cells after 3 h and 6 h in TB medium at 37 °C. n = 1200.

### CapV is a patatin-like phospholipase, which alters the steady-state lipid profile

Blast search of CapV_ECOR31 showed that CapV homologs with >60% identity are not only found in individual *E. coli* and *V. cholerae* strains, but are widely distributed among gamma-proteobacterial species including *Yersinia, Salmonella, Pseudomonas, Shewanella* and *Klebsiella* (fig. S6A). Phylogenetic analysis of representative CapV_ECOR31 homologs supported classification into four different subgroups (fig. S6B). Alignment of the amino acid sequences of those CapV homologs showed that Q329 is an invariant amino acid among even distantly related CapV proteins (fig. S6A). As CapV from *V. cholerae* (*42*), CapV_ECOR31 contains an N-terminal canonical patatin-like phospholipase A2 (PNPLA) domain with three main characteristic conserved signature motifs (*50, 51*), the phosphate or anion binding motif G-G-G-x-[K/R]-G, the esterase box G-x-S-x-G, and the D-G-[A/G] motif as part of the catalytic dyad (Fig. 5A and B). The G-x-S-x-G motif includes the conserved nucleophilic serine of the active site characteristic for the phospholipase A_2_ (PLA_2_) superfamily (*51, 52*). To investigate if serine is required for swimming inhibition and filamentation upon CapV_Q329R_ overexpression, a catalytically inactive S64A variant of the protein (CapV_Q329R/S64A_) was generated. Compared with CapV_Q329R_, overexpression of CapV _Q329R/S64A_ equally inhibited swimming motility and induced filamentation (Fig. 1A, Fig. 2D and E), demonstrating that the G-x-S-x-G motif of CapV_Q329R_ is not required for the phenotype. However, substitution of arginine in the G-G-G-x-[K/R]-G motif (CapV_Q329R/R27A_) and aspartic acid of the D-G-[A/G] motif (CapV_Q329R/D197A_) by alanine relieved both the repression of swimming motility and filamentation. Substitution of the two structural glycine residues (CapV_Q329R/G24A/G25A_) of the G-G-G-x-[K/R]-G still partially suppressed swimming motility and induced a mild filamentous phenotype upon overexpression of the protein (Fig. 1A and B, Fig. 2D and E). In summary, the G-G-G-x-[K/R]-G and D-G-[A/G] motifs of CapV_Q329R_ are required for repression of the swimming phenotype and cell filamentation.

**Fig. 5.**
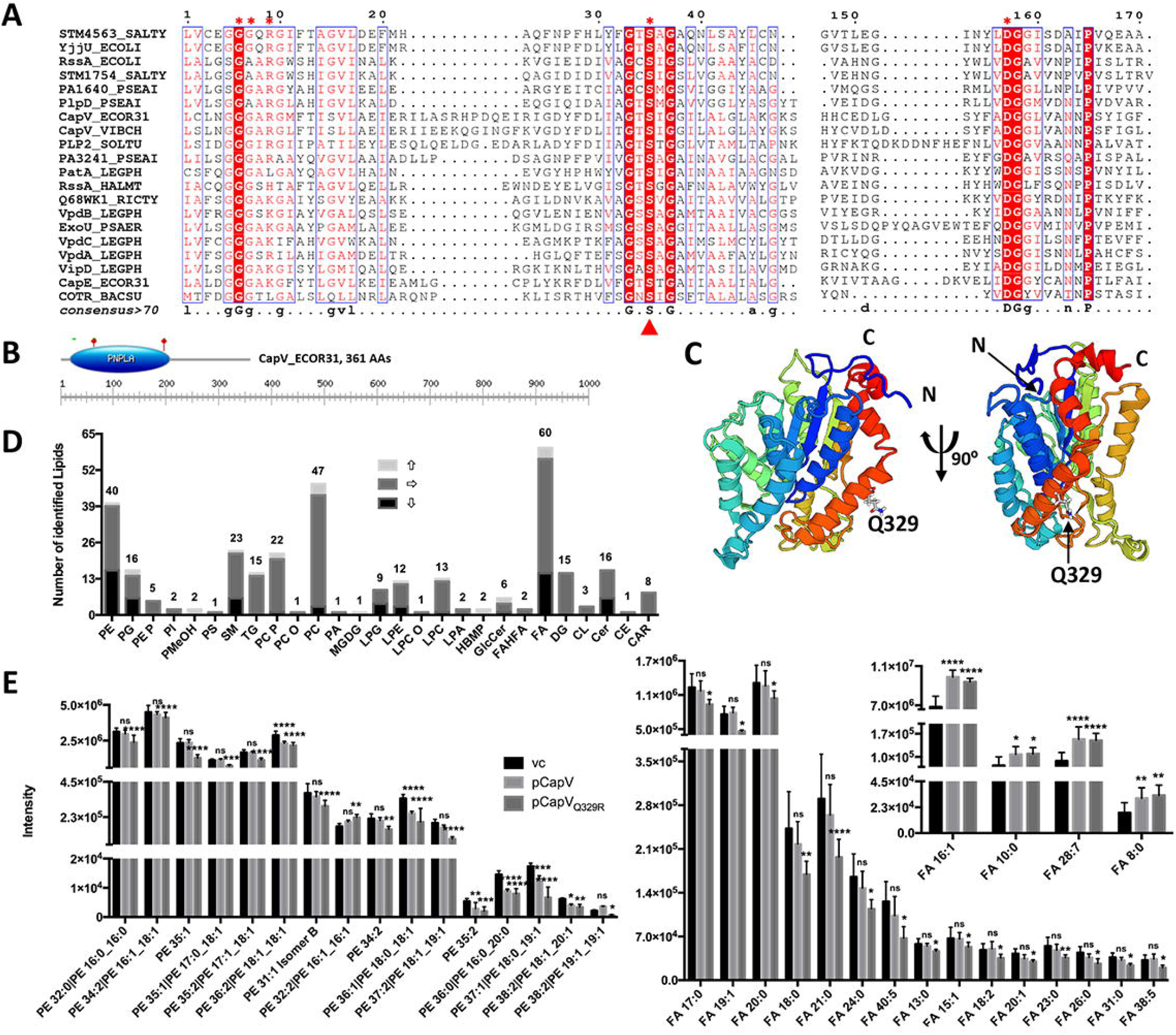
Bioinformatic analysis of the patatin-like phospholipase CapV of *E. coli* MG1655. (**A**) Sequence alignment of CapV_ECOR31 and selected known phospholipase from other species establishes the conserved motifs of the PNPLA domain. Entirely conserved residues are shown in white on a red background. Conserved residues are boxed. The putative catalytic residues of CapV are indicated with filled red triangles. The residues in CapV_Q329R_ mutated to alanine are marked with red asterisks above the sequence. The consensus sequence at the bottom indicates in uppercase letter residues with 100% identity and in lowercase letter residues with higher than 70% conservation. Alignment was performed using CLUSTALW (81), and the result was processed with ESPript 3.0 (*82*). Sequences are: STM4563_SALTY: *S. typhimurium* NP_463419.1, YjjU_ECOLI: *E. coli* BAE78366.1, RssA_ECOLI: *E. coli* NP_415750.2, STM1754_SALTY: *S. typhimurium* YchK NC_003197.2, PA1640_PSEAI: *Pseudomonas aeruginosa* SG17M Homolog_EWH27047.1, CapV_ECOR31: *E. coli* ECOR31 OII97420.1, CapV_VIBCH: *V. cholerae* NC_002505.1, PLP2_SOLTU: *Solanum tuberosum,* PA3241_PSEAI: *P. aeruginosa* SG17M Homolog_EWH29020.1, PatA_LEGPH: *Legionella* lpg2317 NC_002942.5, RssA_HALMT: *Haloferax mediterranei* WP_004060664.1, Q68WK1_RICTY: *Rickettsia typhi* RT0522 WP_011190972.1, VpdB_LEGPH: *Legionella pneumophila* QGK66366.1, ExoU_PSAER: *P. aeruginosa* WP_003134060.1, VpdC_LEGPH: *L. pneumophila* QGK66518.1, VpdA_LEGPH: *L. pneumophila* CCD09784.1, VipD_LEGPH: *L. pneumophila* WP_010948518.1, CapE_ECOR31: *E. coli* ECOR31 OII97423.1, COTR_BACSU: *Bacillus subtilis* WP_003243674.1. (**B**) The graphical representation and schematic indication of the positions of the conserved motifs (indicated by green bar) and putative active site (marked by red stars) in the PNPLA domain of CapV_ECOR31. The graph was assessed by ExPASy_Prosite. (**C**) Predicted structural model of the CapV_ECOR31 shown as ribon representation. The structural model was built with the I-TASSER server (*84*), the result was processed with SWISS-MODEL (*85*). The model was prepared using the coordinates of the 22% identical protein FabD from *Solanum cardiophyllum* (PDB: 1oxwC). (**D**) Number and alteration of identified lipid species in CapV_Q329R_ induced filamentous cells compared to *E. coli* MG1655 VC and relative abundance of PE (**E**) and FA (**F**) derivatives by untargeted CSH-QTOF MS analysis. PE, phosphatidylethanolamine; PG, phosphatidylglycerol; PE P, plasmenyl-phosphatidylethanolamine; PI, phosphatidylinositol; PMeOH, phosphatidylmethanol; PS, phosphatidylserine; SM, sphingomyelin; TG, triglyceride; PC P, plasmenyl-phosphatidylcholine; PC O, plasmanyl-phosphatidylcholine; PC, phosphatidylcholine; PA, phosphatidic acid; MGDG, monogalactosyldiacylglycerol; LPG, lysophosphatidylglycerol; LPE, lysophosphatidylethanolamine; LPC O, plasmanyl-lysophosphatidylcholine; LPC, lysophosphatidylcholine; LPA, lysophosphatidic acid; HBMP, 1-Monoacylglycerol-phospho-2,3-diacylglycerol; GlcCer, glycosylceramide; FAHFA, fatty acid ester of hydroxyl fatty acid; FA, fatty acid; DG, diacylglycerol; CL, cardiolipin; Cer, ceramide; CE, cholesteryl ester; SM, sphingomyelin; CAR, acylcarnitine. Bars represent mean values from six independent replicates with error bars to represent SD. Differences between mean values were assessed by two-tailed Student’s t-test (ns, not significant; *p < 0.05, **p < 0.01, and ***p < 0.001 compared to MG1655 VC). VC = pBAD28. pCapV = CapV cloned in pBAD28; pCapV_Q329R_ = CapV_Q329R_ cloned in pBAD28.

As Q329 is a conserved amino acid residue in CapV homologs, predicted not to be required for catalytic activity and not part of described consensus motifs, we wanted to clarify the position and the effect of the Q329R mutation in the protein. Replacement of Q329 by lysine still partially repressed the apparent swimming motility (Fig. 1B). To this end, we generated a structural model of CapV using the closest homolog in the PDB database, the 22% identical lysophospholipase-like protein FabD from *Solanum cardiophyllum* (PDB: 1oxwC; Fig. 5C), as a template. According to this model, Q329_CapV is located within helix 12 the second last α helix in the context of the RARGRR_329_ sequence pointing outward with no change in the overall structure of the monomer or potential oligomer assembly.

In order to assess whether overexpression of CapV and CapV_Q329R_ caused significant changes in the lipid profile concomitant with filamentation, we extracted lipids and subjected the extracts to mass spectrometry. The lipids, known to be most abundant in the E. coli membrane, phosphatidylethanolamines (PE), phosphatidylglycerols (PG (and cardiolipin (CL) displayed the highest peak intensity. We subsequently observed significant changes in the peak intensity of members of most abundant phospholipid classes such as phosphatidylethanolamines (PE) and phosphatidylglycerols (PG), but also of free fatty acids, lysophospholipids, phosphatidylcholines (PC), ceramides and sphinomyelins, although the peak intensity of the latter three classes was at least 100-fold lower (Fig. 5D,E,F and fig. S7).

### Vitamin B6 restricts cell filamentation induced by CapV_Q329R_

Filamentation has been shown to be affected by environmental conditions (*23*). We found that filamentation was particularly pronounced during growth in TB medium (1% tryptone, 0.5% NaCl), while it was restricted in LB medium (1% tryptone, 1% NaCl, 0.5% yeast extract). Supplementation of TB medium with 0.5% yeast extract (YE) repressed the filamentous phenotype dramatically, while 5% restored the rod-shape of all cells. Supplementation with 0.5% NaCl did not affect filamentation (Fig. 6A).

**Fig. 6.**
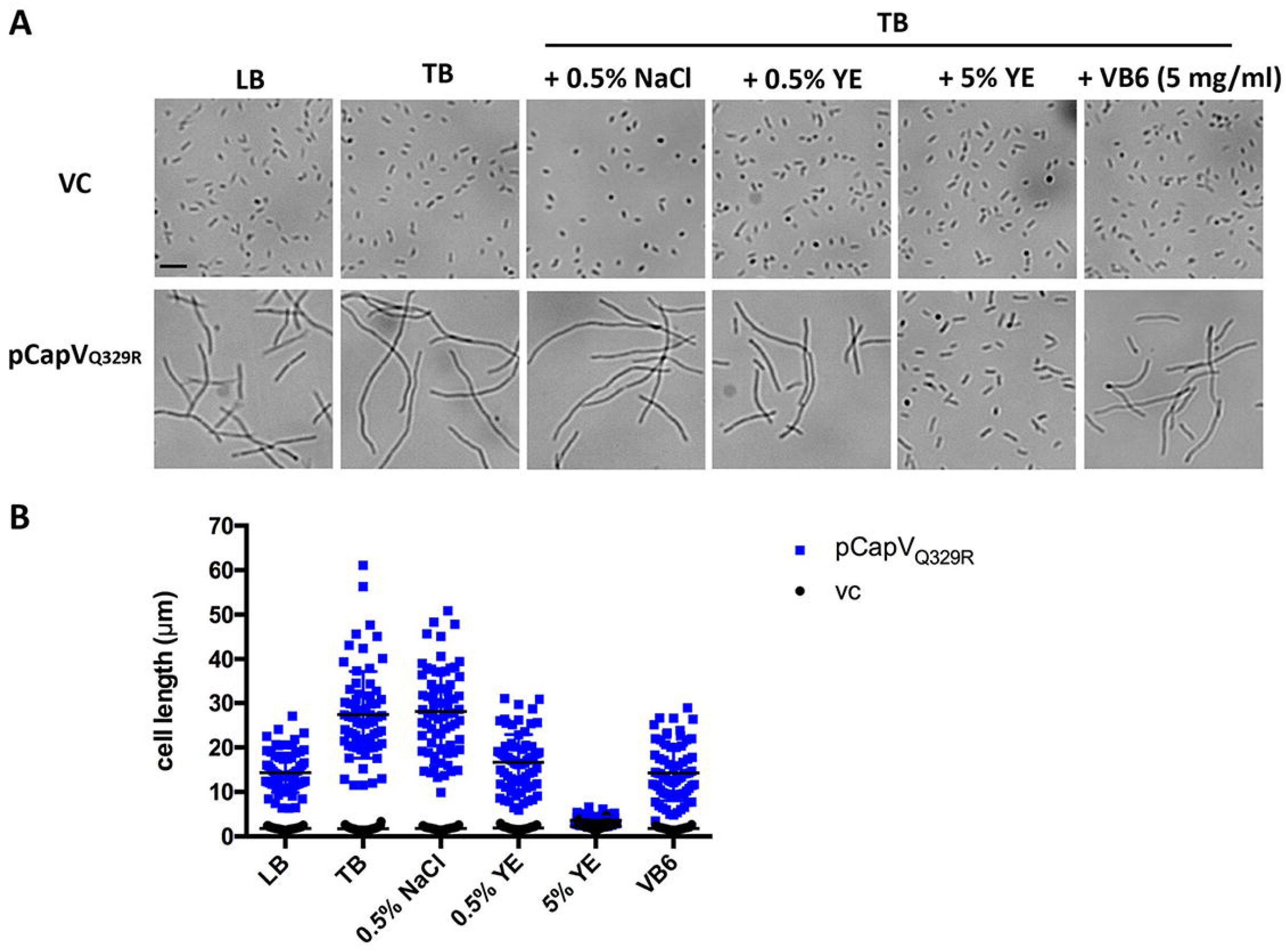
Vitamin B6 (pyridoxine) restricts cell filamentation of *E. coli* MG1655 induced by CapV_Q329R_. (**A** and **B**) Light microscopy pictures of *E. coli* MG1655 cell morphology and quantification of cell length in LB and TB supplemented with 0.5% NaCl, 0.5% YE, 5% YE, and VB6 (pyridoxine, 5 mg/ml), respectively. The quantification is based on results from at least three independent experiments with assessment of 70 cells from each group. Bar, 5 μm. VC = pBAD28.

Yeast extract is the water-soluble portion of autolyzed yeast cells and usually used to prepare microbiological culture media for bacterial studies (*53*). As a nutrient source, it provides nitrogen, amino acids, peptides, carbohydrates, vitamin B complex and other components that promote microbial growth (*54*). In particular, yeast extract contains B-vitamins, the water-soluble precursors of enzyme cofactors with unrelated structure: thiamine (B1), riboflavin (B2), nicotinamide (B3), pantothenate (B5), pyridoxine (B6), biotin (B7), folic acid (B9), and cobalamin (B12) belong to the B-vitamin complex. To identify the component in YE that contributes to the repression of cell filamentation upon CapV_Q329R_ overexpression, we supplemented TB medium with individual B vitamins. While supplementation with thiamine, riboflavin, nicotinamide, pantothenate, biotin, folic acid, and cobalamin showed no effect, supplementation with pyridoxine (B6) at 5 mg/ml decreased the length of filaments by 50% (Fig. 6A and B, fig. S8). To our knowledge, this is the first report about pyridoxine to inhibit of bacterial cell filamentation.

### The effect of CapV_Q329R_ extends to commensal and UPEC *E. coli* strains and *S. typhimurium*

To determine if the effect of CapV_Q329R_ is restricted to *E. coli* K-12 MG1655, we overexpressed CapV_Q329R_ in commensal and UPEC *E. coli* strains (*55, 56*) and *S*. *typhimurium* UMR1. In all cases, CapV_Q329R_ overexpression inhibited apparent swimming motility (fig. S9A and B).

Furthermore, CapV _Q329R_ induced different degrees of cell filamentation in *E. coli* strains and *S*. *typhimurium* UMR1 (fig. S9C). In *E. coli* Fec32 and Fec89, CapV_Q329R_ expression induced mild cell filamentation, while moderate cell filamentation was observed in Fec67 and extensive filamentation in ECOR31, CFT073, and *S*. *typhimurium* UMR1. Interestingly, moderately filamented cells as well as individual rod-shaped cells were observed in the UPEC strain *E. coli* No. 12 and the commensal *E. coli* strain Tob1 upon CapV_Q329R_ overexpression (fig. S9C).

### Expression of CapV_Q329R_ affect rdar biofilm formation on agar plates

We were subsequently wondering, whether CapV_Q329R_ affects phenotypes other than cell filamentation and flagella expression. To this end, we investigated the rdar (red, dry, and rough) biofilm morphotype on agar plates. Although *E. coli* MG1655 and CFT073 displayed a saw (smooth and white) morphotype on LB without salt agar plate after 24 h at 37 °C upon CapV_Q329R_ overexpression, the phenotype was altered after 48 h (fig. S9D). In the same line, *S*. *typhimurium* strain UMR1 displays saw after 24 h (*57*), with its morphology to be dramatically changed by CapV_Q329R_ overexpression after 48 h at 37 °C (fig. S9D).

We then investigated strains that displayed the rdar phenotype at 37 °C. We choose the UPEC strain No. 12, which expresses a semi-constitutive *csgD*-dependent rdar morphotype (*56*). While overexpression of CapV wild type had no effect on rdar phenotype expression compared to No. 12 VC, overexpression of CapV_Q329R_ dramatically decreased the rdar phenotype (Fig. 7A). Scanning and transmission electron microscopy (TEM) of colonies showed that CapV_Q329R_ induced even more extensive cell filamentation on agar plates compared to liquid culture, without affecting cell arrangements (Fig. 7B, fig. S9C and data not shown). While nonfilamented cells produced a pronounced extracellular matrix, the filaments produced only little or no matrix (Fig. 7B).

**Fig. 7.**
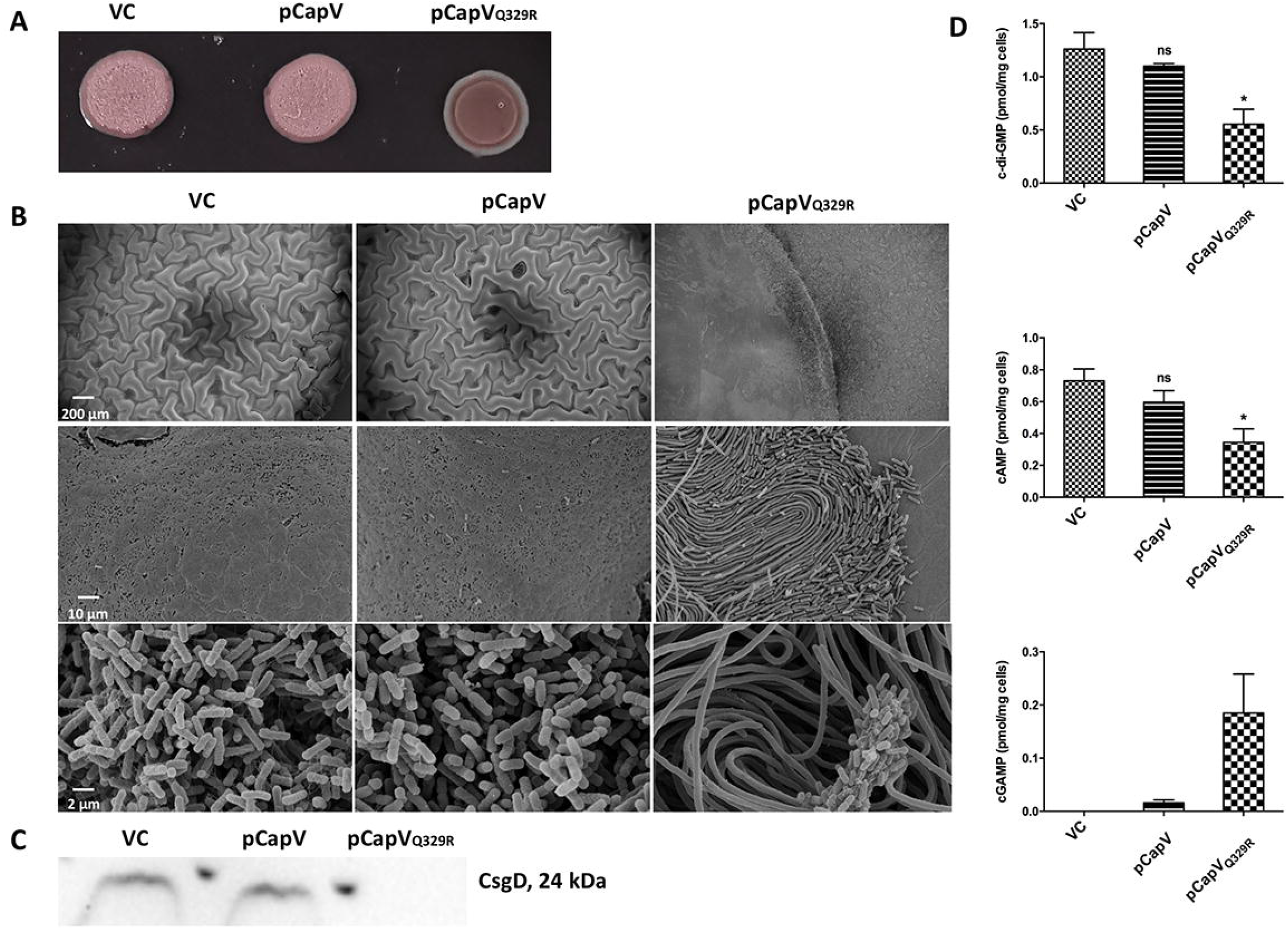
CapV_Q329R_ expression modulates rdar biofilm formation, CsgD expression, and cyclic (di)nucleotides levels in *E. coli* strain No. 12. (**A**) Rdar morphotype in wild-type *E. coli* No. 12 and upon overexpression of CapV and its mutant CapV_Q329R_. Cells were grown on a salt-free LB agar plate for 24 h at 37 °C. VC = pBAD28. (**B**) Scanning electron microscopy of cells from plate-grown colonies. Cells were harvested from a salt-free LB agar plate after 24 h of growth at 37 °C. (**C**) CsgD production upon overexpression of CapV_Q329R_. Only colony morphotypes from the same plate and signals from the same Western blot are compared. (**D**) LC-MS/MS quantification of *in vivo* amounts of c-di-GMP, cAMP, and cGAMP upon overexpression of CapV_Q329R_. Data are displayed as absolute amounts referred to the original cell suspension.

In conjunction with downregulation of the rdar morphotype, the rdar morphotype promoting transcriptional regulator CsgD was downregulated (Fig. 7C), demonstrating that CapV_Q329R_ expression affects the central hub of rdar biofilm formation. Furthermore, CapV_Q329R_ overexpression downregulated the rdar morphotype in the commensal *E. coli* strains ECOR31, Fec67, Fec89, and Tob1 (fig. S9D).

### CapV_Q329R_ expression alters cyclic (di) nucleotide concentrations

In *Enterobacteriaceae*, cyclic di-GMP is a ubiquitous bacterial second messenger, which stimulates the rdar biofilm morphotype via *csgD* expression (*36, 58, 59*). We investigated the cyclic (di) nucleotide levels exemplarily in the UPEC strain *E. coli* No. 12. Along with inhibition of *csgD* expression, the *in vivo* cyclic di-GMP level was concomitantly decreased upon CapV_Q329R_ overexpression (Fig. 7D). Besides cyclic di-GMP, both cAMP (*60*) and cGAMP (*39*) regulate *E. coli* biofilm formation. Consistent with a downregulated rdar biofilm phenotype, a reduction in cAMP and an increase in cGAMP levels was observed upon CapV_Q329R_ overexpression (Fig. 7D). Taken together, these results indicate CapV_Q329R_ inhibits rdar biofilm formation potentially by regulating various cyclic (di)-nucleotide signals.

Taken together, our results showed that the observed phenotypes upon CapV_Q329R_ expression were not specific to *E. coli* MG1655, but were common to other commensal and UPEC *E. coli* strains and *S. typhimurium* UMR1, indicating a general effect of CapV_Q329R_ on bacterial cell morphology, regulation of swimming motility and biofilm formation.

### CapV_Q329R_ enhances susceptibility of *E. coli* MG1655 to bacteriophage P1 infection and the antibiotic cephalexin

Recently, the patatin-like phospholipase CapV and the CD-NTase DncV has been shown to take part in bacterial antiphage defence systems (*61*). Since CapV_Q329R_ shows a physiological function independent of DncV, we wondered whether CapV_Q329R_ still contributes to antiphage defence. While overexpression of CapV wild type has no effect, interestingly, an approximately 10-fold higher plaque formation was observed upon CapV_Q329R_ overexpression compared to the vector control when MG1655 cells were infected with P1 phage (fig. S10A and B), indicating that CapV_Q329R_ enhances susceptibility to bacteriophage P1 infection.

We also observed that overexpression of CapV_Q329R_ makes *E. coli* MG1655 more susceptible to the cell wall inhibiting antibiotic cephalexin (data not shown). However, a systematic investigation of altered susceptibility to various antibiotic classes awaits to be performed.

## Discussion

Rod-shaped bacteria can undergo filamentation upon dysregulation of cytokinesis components. In this study, we describe a mutant of the patatin-like phospholipase CapV containing a single amino acid change, CapV_Q329R_, to repress the apparent swimming motility, but to induce extensive *sulA*-independent cell filamentation, while only slightly decreasing FtsZ levels in *E. coli* MG1655. Besides these dramatic morphological changes, which occur in commensal and pathogenic *E. coli* strains, modulation of rdar biofilm formation occurs concomitantly. Alteration in the concentrations of cyclic (di)-nucleotides upon CapV_Q329R_ overexpression might be involved in motility and rdar biofilm regulation. Collectively, these findings demonstrate a single amino acid change to create a protein, which dramatically modulates various aspects of bacterial physiology.

Bioinformatic analysis confirmed CapV_ECOR31 is a patatin-like PLA2 (PLP) that contains a PNPLA domain (Fig. 5A and B). PLP proteins are predominantly phospholipase A2 enzymes, acyl hydrolases cleaving the sn-2 position of phospholipids. We did not observe fundamental alterations in the lipid profile concomitant with extensive filamentation after four hours of CapV_Q329R_ induction indicating that CapV and CapV_Q329R_ shows only a minor, if any, catalytic activity in the genetic background of *E. coli* MG1655 (Fig. 5 D,E,F; fig. S7). In plants, PLPs do not only act as enzymes to cleave fatty acids from membrane lipids, but also aid in the control the spread of infection. Such a protein is, for example, highly abundant in potato tuber storage (*62, 63*). In mammals, PLPs are mostly involved in lipid metabolism and turnover (*64*), with a polymorphism in a patatin phospholipase to provide the genetic predisposition for nonalcoholic fatty liver disease and metabolic syndrome (*65*). PLPs are also found in many pathogenic bacterial species and act as toxins in host-pathogen interactions (*66, 67*). One of the best characterized PLPs is ExoU, a cytotoxic effector protein of *P. aeruginosa* secreted through the type III secretion system upon host cell contact (*68*). Host ubiquitination triggers PLA2 activity of ExoU leading to degradation of host cell membranes (*69*). Our finding that the G-G-G-x-[K/R]-G and D-G-[A/G] motifs in the PNPLA domain of CapV_Q329R_, but not the catalytic G-x-S-x-G motif, are required for the cell filamentation phenotype suggests a functional role for the protein scaffold (Fig. 2D). Since bacterial filamentation also contributes to pathogenesis by escape from phagocytosis, we hypothesize that CapV_Q329R_ has a phenotype in bacterial survival upon induction of cell filamentation during interaction with host immune cells.

The amino acid Q329 is located outside of the putative PNPLA domain of CapV_ECOR31 and consequently not part of characteristic motifs required for the catalytic activity of this PLA2 superfamily member (Fig. 5B). 3D model construction using the closest homolog of CapV_ECOR31 in the PDB database (FabD from *Solanum cardiophyllum,* PDB: 1oxwC) revealed that Q329_CapV is located in the context of the RARGRR_329_ sequence motif within the second last α helix of the CapV structural model with the arginine side chain pointing outwards causing no change in the overall structure of CapV_ECOR31 (Fig. 5C). However, the switch from a potential hydrogen-bond acceptor (Q) to a hydrogen-bond donor (R) with a longer side chain indicates that the CapV_ECOR31 mutant might show altered enzymatic activity, ligand binding properties or protein-protein interactions. Arginine with its positively charged side chain can be involved in a variety of different functionalities such as in binding of negative charged phosphates molecules in DNA molecules or nucleotides. The RR twin arginine motif is part of the N-terminal signal sequence for the Twin-Arginine Translocation (TAT) pathway (*70, 71*). Furthermore, a RxxR motif constitutes a conserved peptidase cleavage site, while a RxxxR motif is part of the binding motif for c-di-GMP in PilZ domain proteins, whereby the arginine residues can bind O-6 and N-7 at the Hoogsteen edge of the guanine base (*72*).

The catalytic activity of bacterial and eukaryotic patatin-like phospholipases is highly regulated to require a cofactor for activation (*51*). CapV from *V. cholerae* El Tor is activated by binding to 3’3’-cGAMP synthesized by the cyclic dinucleotide cyclase DncV to cause growth retardation (*42*). Up to now, binding site of 3’ 3’-cGAMP in CapV has not been identified. Whether variant CapV_Q329R_ shows altered 3’ 3’-cGAMP binding needs to be tested. Our result showed that the *E. coli* homolog CapV_ECOR31 variant CapV_Q329R_ did hardly affect MG1655 cell viability at logarithmic phase growth, but induced a mild 50% growth retardation in stationary phase under our experimental conditions (Fig. 4, fig. S5). We cannot exclude 3’ 3’-cGAMP binding to be required for this or other phenotypes. However, the *in vivo* 3’3’-cGAMP signal was under the detection limit in the *E. coli* K-12 strain (data not shown). As the catalytic serine residue in the G-x-S-x-G motif of the PNPLA domain is not involved in the activation of cell filamentation, we cannot exclude that CapV_Q329R_ might induce cell filamentation solely by its protein scaffold (independent of 3’ 3’-cGAMP binding) or an enzymatic activity other than a phospholipase activity.

Bacterial cell filamentation is often associated with activation of the SOS response and the cell division inhibitor SulA (*14, 29*). According to our result, CapV_Q329R_ induced a *sulA*-independent filamentation in MG1655 (fig. S4D), indicating CapV_Q329R_ affects cell morphology possibly via an alternative signaling pathway.

Besides filamentation, CapV_Q329R_ overexpression inhibited apparent swimming motility of MG1655 in the soft agar plate (Fig. 1A), with CapV_Q329R_ overexpression cells concomitantly to gradually loose flagella to become non-motile (movie S1). Long filamentous cells, even if observed being motile when up to 20 times the length of a wild type cell in liquid culture, might become physically trapped in the pores of the agar.

Interestingly, CapV_Q329R_ also downregulated rdar biofilm formation in several *E. coli* strains that express the morphotype at 37 °C (fig. S9D, Fig. 7A). Examination of cells showed that cells are highly filamented along with decreased rdar biofilm formation (Fig. 7B), suggesting CapV_Q329R_ remodels colony morphology on the agar plate. Besides the reduction of *in vivo* c-di-GMP level, a reduction in cAMP level and a weak signal for 3’ 3’-cGAMP were detected upon CapV_Q329R_ overexpression in the UPEC *E. coli* strain No. 12 (Fig. 7D). Cyclic AMP has been shown to promote extracellular matrix production and biofilm formation in UPEC (*60*). Our previous studies showed that DncV-synthesized 3’3’-cGAMP participates in down-regulation of rdar biofilm formation in *E. coli* ECOR31 (*39*). Besides DncV, a certain class of GGDEF domain proteins has been identified to produce 3’3’-cGAMP (*73*). Inspection of the genome sequence did not indicate the presence of a *dncV* homolog in the *E. coli* No. 12 genome (unpublished data), suggesting that a 3’3’-cGAMP production of a GGDEF domain protein(s) produces might be directly or indirectly activated by CapV_Q329R_, to inhibit rdar biofilm formation of *E. coli* No. 12.

We show in this work that CapV_Q329R_ expression increased susceptibility to infection by the myophage P1 (fig. S10). Of note, the *capV-dncV-vc0180-vc0181* four-gene operon (fig. S1) of *E. coli* and *V. cholerae* confers immunity against various bacteriophages in several gene product combinations (*61*). Basically, upon phage infection, the cyclic di-nucleotide cyclase DncV synthesizes 3’3’-cGAMP which in turn activates the patatin-like phospholipase CapV to lead to cell death and eventually abort infection (*61*).

In conclusion, our study reported CapV_Q329R_, a putative patatin-like phospholipase, to inhibit apparent swimming motility and rdar biofilm formation, while to promote extensive cell filamentation and susceptibility to phage P in *E. coli* MG1655. Several of these physiological effects were shown to extend to various *E. coli* strains and *S. typhimurium,* suggesting a single amino acid change to evolve a protein which has the ability to dramatically modulate various aspects of bacterial physiology.

## Materials and Methods

### Bacterial strains and growth conditions

All strains used in this study are listed in table S1. Strains were cultured in tryptone broth (TB, 1% tryptone, 0.5% NaCl), Luria-Bertani (LB, 1% tryptone, 0.5% yeast extract, 1% NaCl) liquid medium or on LB agar plates supplemented with 25 μg/ml chloramphenicol at indicated temperatures. L-arabinose (Sigma) was used for induction of gene expression at indicated concentrations.

### Plasmid construction

All plasmids and primers used in this study are listed in table S1 and table S2, respectively. Genes of interest were amplified by PCR, the PCR products digested with indicated restriction enzymes and ligated into the pBAD28 vector using the Rapid DNA Ligation Kit (Roche Diagnostics). Inserted DNA sequences were confirmed by DNA sequencing (StarSeq).

### Site-directed mutagenesis

Site-directed mutagenesis was performed using the Q5 site-directed mutagenesis kit according to the manufacturer’s instructions (NEB). All mutations were confirmed by DNA sequencing.

### Swimming assay

To assess apparent swimming motility, 3 μl of an overnight culture suspended in water (OD_600_ = 5) was inoculated into soft agar medium containing 1% tryptone, 0.5% NaCl, and 0.25% agar (*74*). The swimming halo was documented with a Gel Doc imaging system (Bio-rad) after 6 h at 37 °C and the swimming diameter was measured by ImageJ 1.8.0 (*75*).

### Rdar colony morphotype assessment

To visualize expression of cellulose and curli fimbriae, 5 μl of an overnight culture of OD_600_ = 5 suspended in water was spotted onto LB without NaCl agar plates containing the dye Congo red (40 μg/ml, Sigma) and Coomassie Brilliant Blue G-250 (20 μg/ml, Sigma). Plates were incubated at 37 °C. Pictures were taken at different time points to analyze the development of the colony morphology structure and dye binding.

### Isolation of cell-associated flagellin

Bacterial cell-associated flagellin was isolated as described previously (*76*). Briefly, a single bacterial colony of each group was inoculated in TB medium and cultured overnight at 37 °C at 150 rpm. After dilution to OD_600_ = 0.01, 0.1% L-arabinose was added for induction of the protein and culturing was continued for indicated time points. Flagella were sheared off by forcing the sample 15 times through a syringe with a needle of 0.51 mm diameter (BD Microlance). One ml of samples was collected, centrifuged at 17000 rpm and the supernatant was mixed with cold trichloroacetic acid (v:v = 3:1, Sigma). Samples were incubated at −20 °C for 2 h, followed by centrifugation at 17000 rpm for 40 min at 4 °C. The cell pellet was collected for SDS-PAGE analysis (4% stacking gel, 12% running gel) and the gel was stained by colloidal Coomassie brilliant blue (Sigma).

### Flagella staining

Staining of bacterial flagella according to Leifson was performed using a flagella stain kit according to the instructions of the manufacturer (BASO Diagnostics). Flagella were observed under a light microscope (Leica MC170 HD).

### Transmission Electron Microscopy

Bacterial flagella were visualized by transmission electron microscopy (TEM) (*39*). Briefly, a suspension of bacterial cells grown overnight was diluted in TB medium with protein expression induced by 0.1% L-arabinose for indicated time points. An aliquot of 3 μl from each sample was added to a grid with a glow discharged carbon coated supporting film for 3 min. The excess solution was soaked off by a filter paper and the grid was rinsed by adding 5 μl distilled water for 10 sec. Distilled water was soaked off by a filter paper and the grid was stained with 5 μl 1% uranyl acetate (Sigma) in water for 7 sec. Excess stain was soaked off by a filter and the grid air-dried. The samples were examined in a Hitachi HT 7700 (Hitachi, Tokyo, Japan) electron microscope at 80 kV and digital images were taken with a Veleta camera (Olympus, Münster, Germany).

### Scanning electron microscopy

The bacterial cells were fixed in 1.5 ml fixative solution (0.5 % glutaraldehyde, 2.5 % paraformaldehyde in 10 mM HEPES, pH 7.0) for 2 h at 4 °C. The fixed biofilm sample was processed for observation by scanning electron microscopy (SEM) applying dehydration with acetone, critical-point drying and sputters coating with gold/palladium. Samples were examined in a Zeiss Merlin field emission scanning electron microscope at an acceleration voltage of 5 kV with the Everhart-Thornley SE-detector and the inlens SE-detector in a 30:70, 70:30 or 77:23 ratio.

### Light and fluorescence microscopy

Cell morphology was investigated either under the light microscope (Leica MC170 HD) or a fluorescence microscope (Nikon). Viability of cells was analyzed with the LIVE/DEAD BacLight fluorescence stain from Life Technologies (Thermo Fisher Scientific). For all measurements, live cells were mounted onto 1% agarose pads supplemented with TB medium prior to microscopy analysis. Phase-contrast and fluorescence images were captured using a Ti eclipse inverted research microscope (Nikon) with a 100x/1.45 numerical aperture (NA) objective (Nikon). Image processing and cell length measurement was conducted by Fiji ImageJ 1.8.0 (*75*).

### Lipid extraction

Lipid extraction was performed as described previously with slight modifications (*42, 77*). Briefly, overnight cultures were inoculated into 50 ml of TB to OD_600_ = 0.01, expression of gene products induced by 0.1% arabinose and grown for 4 h at 37 °C. Cells were collected by centrifugation at 4 °C for 20 min, and 5 ml organic extraction buffer (methanol:chloroform:0.1 M formic acid = 20:10:1, v/v/v) was added to the cell pellets, followed by 1 h of heavy shaking at room temperature. 2.5 ml inorganic aqueous buffer (0.2 M H_3_PO_4_, 1 M KCl) was added, followed by another 1 h of heavy shaking. The mixture was centrifuged at 13000 g for 10 min, lipids dissolved in the lower chloroform phase were collected for analysis by mass spectrometry.

### Lipidomics analysis

Extracted lipids were analyzed using ultra high-pressure liquid chromatography (UHPLC) on a Waters CSH column, interfaced to a quadrupole/time-of-flight (QTOF) mass spectrometer (high resolution, accurate mass), with a 15 min total run time. The LC/QTOFMS analyses are performed using an Agilent 1290 Infinity LC system (G4220A binary pump, G4226A autosampler, and G1316C Column Thermostat) coupled to either an Agilent 6530 (positive ion mode) or an Agilent 6550 mass spectrometer equipped with an ion funnel (iFunnel) (negative ion mode). Lipids are separated on an Acquity UPLC CSH C18 column (100 x 2.1 mm; 1.7 μm) maintained at 65°C at a flow-rate of 0.6 ml/min. Solvent pre-heating (Agilent G1316) was used. The mobile phases consist of 60:40 acetonitrile: water with 10 mM ammonium formate and 0.1% formic acid (A) and 90:10 propan-2-ol: acetonitrile with 10 mM ammonium formate and 0.1% formic acid. The gradient is as follows: 0 min 85% (A); 0–2 min 70% (A); 2–2.5 min 52% (A); 2.5–11 min 18% (A); 11–11.5 min 1% (A); 11.5–12 min 1% (A); 12–12.1 min 85% (A); 12.1–15 min 85% (A).

Samples are injected (1.7 μl in positive mode and 5 μl in negative ion mode) with a needle wash for 20 seconds (wash solvent is isopropanol). The valve is switched back and forth during the run for washing to reduce carryover of less polar lipids. Sample temperature is maintained at 4°C in the autosampler. Data are collected in both positive and negative ion mode, and analyzed using MassHunter (Agilent). Lipids are identified based on their unique MS/MS fragmentation patterns using in-house software, Lipidblast.

### Extraction of *in vivo* produced nucleotides

Extraction of cyclic dinucleotides from bacterial cells was performed as reported (*78*). Briefly, individual colonies were inoculated in TB medium, overnight cultures diluted to OD_600_ = 0.01 and grown in TB medium at 37 °C for 4 h containing 0.1% L-arabinose to induce expression of CapV and CapV_Q329R_. 5 ml cell suspension was pelleted and resuspended in 500 μl ice cold extraction solvent (acetonitrile/methanol/water/formic acid=2/2/1/0.02, v/v/v/v), followed by boiling for 10 min at 95 °C. Three subsequent extracts were combined and frozen at −20 °C overnight. The extracts were centrifuged for 10 min at 20,800 g, evaporated to dryness in a Speed-Vac (Savant) and analyzed by LC-MS/MS.

### Western blot analysis

To detect protein expression, 5 mg (wet weight) of bacterial cells collected from agar plates or liquid cultures were resuspended in 200 μl SDS sample buffer and heated at 95 °C for 10 min. The total protein content was assessed by Coomassie Brilliant blue staining after sample separation. Samples containing equal amounts of protein were separated by SDS-PAGE (4% stacking and 12% resolving gel) and transferred onto a PVDF membrane (Millipore). The membrane was blocked with 5% skim milk (for detection of *E. coli* CsgD and flagellin FliC) or 5% BSA (for detection of His-tagged protein) in blocking buffer overnight. CsgD and FliC were detected with a polyclonal *E. coli* anti-CsgD peptide primary antibody (dilution 1:5000) (*79*) and anti-Flagellin (FliC) antibody (Abcam, dilution 1:5000), respectively. The horseradish peroxidase-conjugated goat anti-rabbit IgG was the second antibody (dilution 1:2000; Jackson ImmunoResearch Laboratories Inc.). An anti·His antibody conjugated to horseradish peroxidase (Penta·His HRP Conjugate, Qiagen) was used to detect 6xHis-tagged proteins. Antibody binding was visualized with ECL light detection reagent (Roche) using Luminescent Image Analyzer (LAS-1000plus, Fujifilm).

### Phage P1 plaque formation assays

The phage P1 plaque formation assay was performed according to previously described methods with slight modifications (*61, 80*). Individual bacterial colonies from overnight cultures were inoculated into TB medium and grown to OD600 = 0.1 at 37 °C, followed by further growth with 0.1% L-arabinose and Cm for another 3 h to induced the expression of CapV and CapV_Q329R_. For the determination of phage infectivity by plaque formation, 500 μl of cells were thoroughly mixed with 4.5 mL modified MMB agar (TB with 0.1 mM MnCl_2_, 5 mM MgCl_2_, 0.35% agar), immediately poured onto Petri dishes containing 20 ml MMB Agar (1.6%) and allowed to cool for 10 min at room temperature. Tenfold serial dilutions of T1 phage lysate in SM Buffer (100 mM NaCl, 8 mM MgSO_4_, 50 mM Tris-HCl pH 7.5) were dropped on top of the double layer agar plate and allowed to dry for 20 min at room temperature. Plates were incubated at 37 °C for 18 h and plaques were counted to compare efficiency of plating.

### Bioinformatic analyses

A BLAST search against the NCBI protein database was performed with standard parameters using the *E. coli* ECOR31 CapV sequence as a query. All distinct protein sequences with >40% identity from different species were selected. Sequences were aligned using CLUSTALW (*81*) and processed with ESPript 3.0 (*82*) using standard parameters. The phylogenetic tree was reconstructed calculating sequence similarity by Maximum Likelihood (ML) in MEGA 7.0 (*83*). A CapV and CapV_Q329R_ structural model was built with the I-TASSER server (*84*) and processed with SWISS-MODEL (*85*).

## H2: Supplementary Materials

### Supplementary Results

Fig. S1. Overexpression of the four gene operon *capV-dncV-vc0180-vc0181* cloned in pBAD28 (p*78901*) downregulates swimming motility and FliC production of *E. coli* MG1655.

Fig. S2. CapV_Q329R__ECOR31, but not other gene products encoded by *p78901* downregulate apparent swimming motility of MG1655.

Fig. S3. CapV_Q329_ production is critical for induction of apparent inhibition of swimming motility in *E. coli* MG1655 by *p78901.*

Fig. S4. CapV_Q329R_-induced cell filamentation is independent of *sulA* and does hardly affect FtsZ and FtsA production.

Fig. S5. Cell viability assay upon production of CapV and CapV_Q329R_ by the LIVE/DEAD™ BacLight™ Viability Kit.

Fig. S6. Phylogenetic and bioinformatic analysis of CapV homologs.

Fig. S.7: Lipidomic analysis of E. coli MG1655 vector control pBAD28 and overexpressing CapV and CapV_Q329R_

Fig. S8. Effects of various B-vitamins on CapV_Q329R_-induced cell filamentation of *E. coli* MG1655.

Fig. S9. Apparent swimming motility (A, B), cell filamentation (C) and rdar biofilm formation (D) of *E. coli* strains Fec32, Fec67, Fec89, No.12, ECOR31, Tob1, CFT073, and *S. typhimurium* UMR1 upon overexpression of CapV_Q329R_.

Fig. S10. Plaque formation of bacteriophage P1 on *E. coli* MG1655 vector control and strains overexpressing CapV and CapV_Q329R_. Vector control=pBAD28.

### Supplementary Tables

Table S1. Bacterial strains and plasmids used in this study.

Table S2. Primer used in this study

### Supplementary Movies

Movie S1. *E. coli* MG1655 cells examined under a light microscope 4 and 6 h after induction of production of CapV wild type and CapV_Q329R_ by 0.1% L-arabinose in TB medium at 37 °C.

Movie S2. *E. coli* MG1655 cells examined under a light microscope 22 h after induction of CapV_Q329R_ production by 0.2% L-arabinose in TB medium at 37 °C.

## General

We thank the following individuals for providing strains: Prof. Ken Gerdes (Newcastle University), Prof. Van Melderen (Université Libre de Bruxelles), Prof. Daniel O. Daley (Stockholm University), Prof. David Sherrat (University of Oxford) and Prof. Sun Nyunt Wai (Umeå University). We thank Prof. Jan-Willem de Gier (Stockholm University) for providing phage P1. We are also grateful to Prof. Kristina Jonas (Stockholm University) for sharing access to the fluorescence microscope and Dr. Kristen Schroeder and Dr. Sulman Shafeeq for support; Aysha Qureshi and Elsa Malmström for experimental support during their internship, and the EM core facility at Karolinska Institutet for processing our samples. Further, we appreciate the help of Ina Schleicher in assisting in preparation of the scanning electron microscopic samples. We also appreciate the support of Prof. Oliver Fiehn and his team at the University of California in Davis in expert lipidomic analyses.

## Funding

F.L. has received a scholarship from the China Scholarship Council (CSC). This work was supported by a grant from the Swedish Research Council for Natural Sciences and Engineering (621-2013-4809) to U.R.

## Author contributions

F.L., and U.R. conceived the study and designed the experiments. F.L., H.B. and M.R. performed the experiments. F.L., M.R. and U.R. analyzed the data. F.L. and U.R. wrote the paper. All authors reviewed and approved the manuscript.

## Competing interests

The authors declare that they have no competing interests.

## Data and materials availability

All data needed to evaluate the conclusions in the paper are present in the paper and/or the Supplementary Materials.

## Notes

### Competing Interest Statement

The authors have declared no competing interest.

