## Supplemental Material for "Morphological and physiological effects of a single amino acid substitution in the patatin-like phospholipase CapV in *Escherichia coli*"

Supplementary Materials

**Supplementary results**

The first ORF in the four-gene operon *capV-dncV-vc0180-vc0181* is *capV* (fig. S1A). CapV possesses a N-terminal patatin-like phospholipase A2 (PNPLA) domain, the enzymatic activity of which is stimulated by 3′3′-cGAMP in *V. cholerae* (*1*). *VC0180* encodes an eukaryotic-like ubiquitin ligase with ThiF (E1)/ E2 domains and *VC0181*, an isopeptidase with a JAB domain, respectively (*1, 2*) (fig. S2A). We had previously cloned the four genes in pBAD28 under the regulation of the pBAD promoter (*3*). To clarify which gene in the putative four-gene operon contributes to inhibition of *E. coli* MG1655 swimming motility, we constructed amino acid substitutions in the catalytic motif of each of the gene products. Expression of the four genes from p*78901* (Table S1) with an amino acid substitution in the D-G-[A/G] motif of CapV (78901_D197A_) did not suppress apparent swimming motility, while substitutions in residues involved in the catalysis of other proteins maintained the suppressive effect (fig. S2 B, C, D and E).

Surprisingly, both overexpression of chromosomal *capV* and *dncV*-*vc0180*-*vc0181* (p*79801*) had no effect on apparent swimming motility of *E. coli* MG1655 (fig. S3A). Sequence analysis showed the presence of several amino acid mutations in the insert of p*78901*, whereby nucleotide alternation in *capV* led to glutamine 329 to be substituted by arginine (Q329R) (fig. S3A). To investigate the molecular basis of apparent swimming repression of p*78901*, we constructed a plasmid that expressed the CapV_Q329R_ variant, which suppressed apparent swimming motility. Moreover, only reversion of the Q329R mutation in the four gene construct resulting in p*78901*_R329Q_ relieved swimming repression, while substitutions in other genes failed to relieve swimming repression (fig. S3B and C).

**Supplementary Figures**

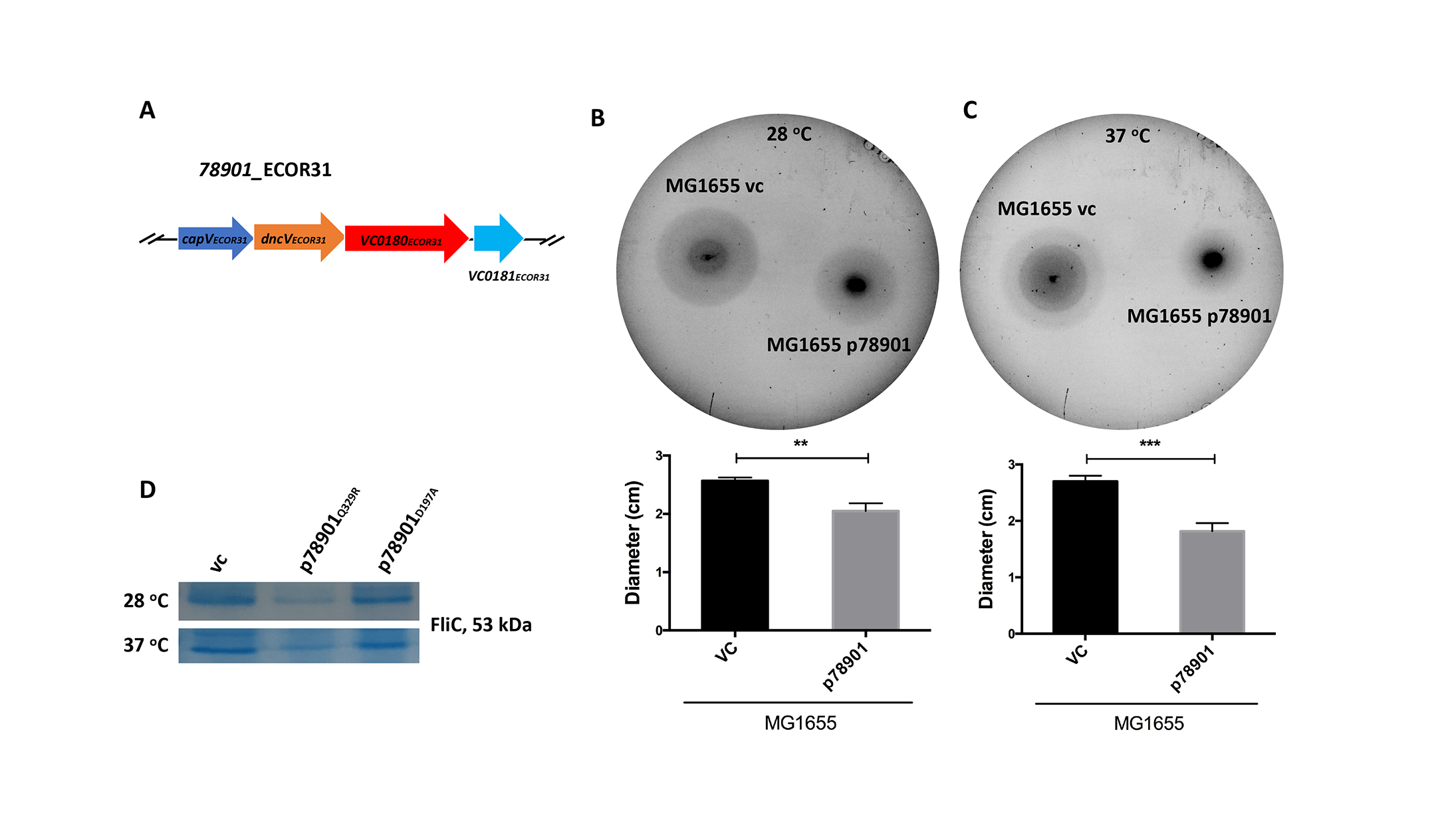

Fig. S1. Overexpression of the four gene operon *capV-dncV-vc0180-vc0181* (A) cloned in pBAD28 (p*78901*) downregulates swimming motility (B and C) and FliC production (D) of *E. coli* MG1655. 3 µl of OD_600_ = 5 cells were inoculated into soft agar plates containing 1% tryptone, 0.5% NaCl and 0.25% agar and the swimming diameter was measured after 8 h at 28 °C and 6 h at 37 °C. Bars represent mean values with error bars to represent SD from three independent replicates. Differences between mean values were assessed by two-tailed Student’s t-test (ns, not significant; *p < 0.05, **p < 0.01, and ***p < 0.001 compared to MG1655 vector control). VC = pBAD28, p*78901* = *capV-dncV-vc0180-vc0181* cloned in pBAD28.

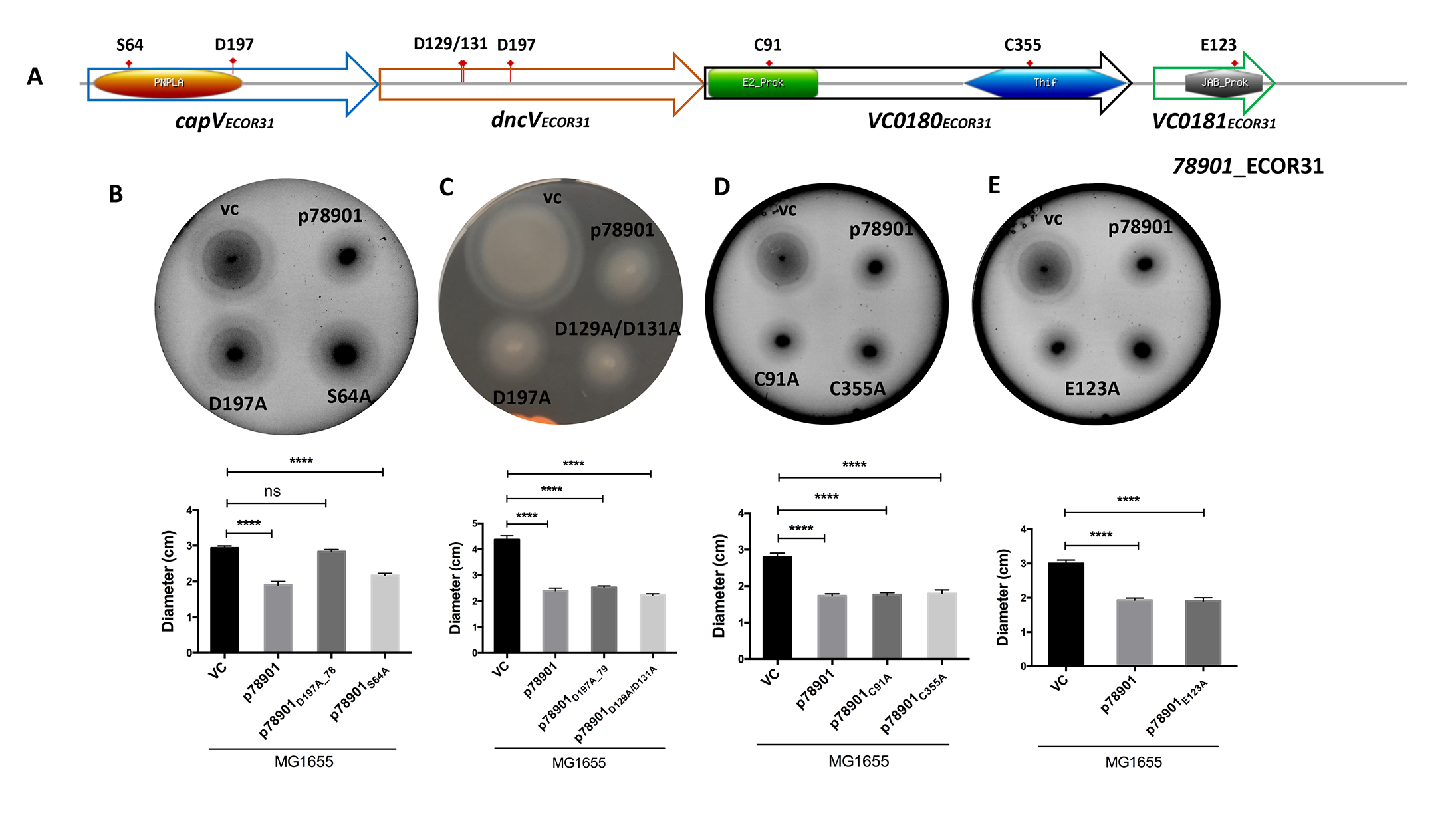

Fig. S2. CapV_Q329R__ECOR31, but not other gene products encoded by p*78901*, downregulates apparent swimming motility of MG1655. (A) Schematic illustration of *E. coli* ECOR31 four gene operon with domain profiles of gene products. The putative catalytic residues are indicated with filled red triangles. Domain prediction was performed with the InterPro server, and the result was processed with ExPASy_Prosite_MyDomains (80). (B-E) Effects of aa substitutions in the catalytic motifs of p78901 encoded gene products on apparent swimming motility of *E. coli* MG1655. 3 µl of OD_600_ = 5 cells were inoculated into soft agar plates containing 1% tryptone, 0.5% NaCl and 0.25% agar and the swimming diameter was measured after 6 h at 37 °C. Bars represent mean values with error bars to represent SD from three independent replicates. Differences between mean values were assessed by two-tailed Student’s t-test (ns, not significant; *p < 0.05, **p < 0.01, and ***p < 0.001 compared to *E. coli* MG1655 vector control). VC = pBAD28, p*78901* = *capV-dncV-vc0180-vc0181* cloned in pBAD28.

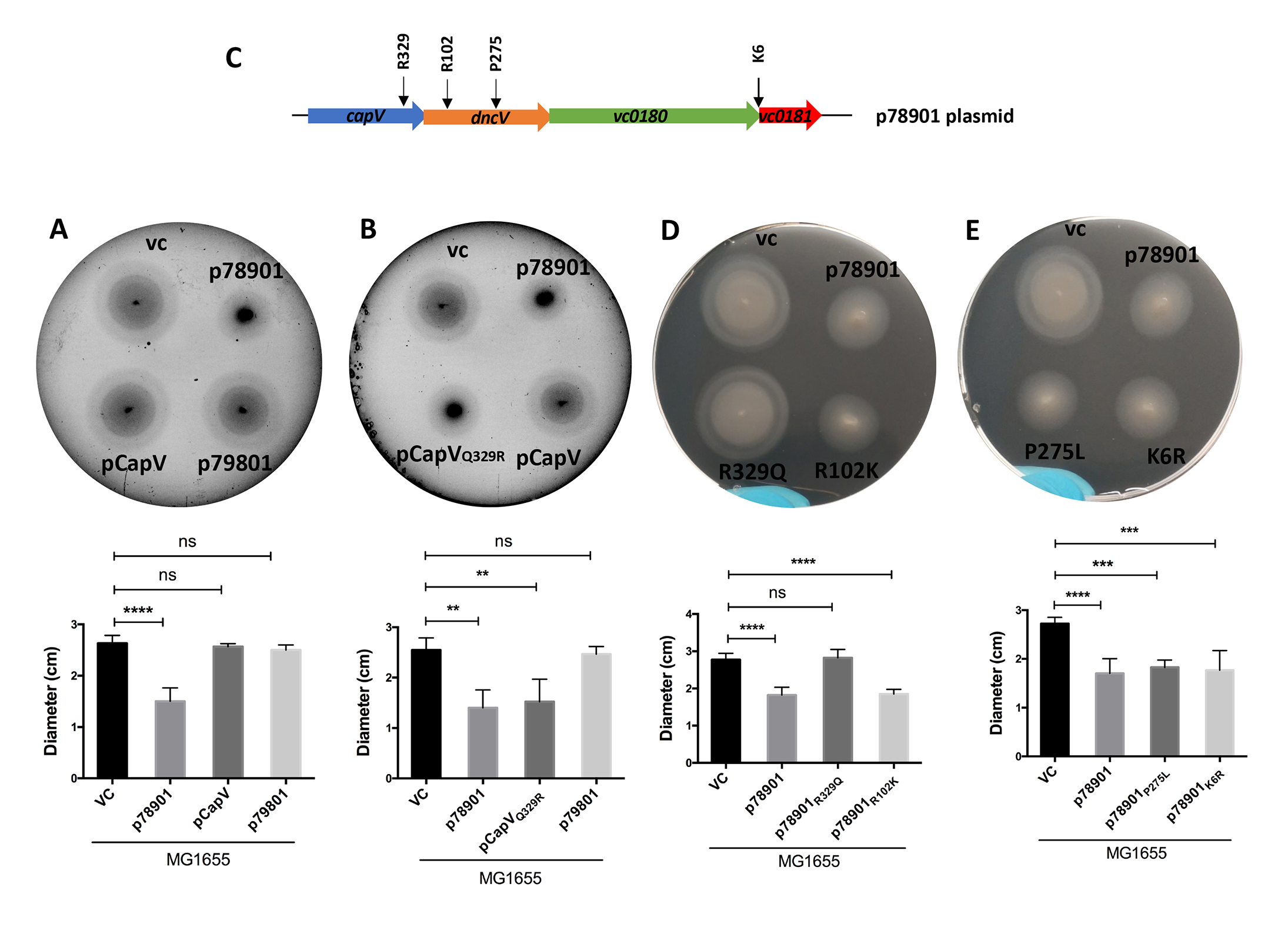

Fig. S3. CapV_Q329_ production is critical for induction of apparent inhibition of swimming motility in *E. coli* MG1655 by p*78901.* (A) Schematic illustration of the aa changes in the p*78901* gene products compared to p*78901* wild type. (B and C) Effects of the aa substitutions in the p*78901* construct on apparent swimming motility of MG1655. 3 µl of OD_600_ = 5 cells were inoculated into soft agar plates containing 1% tryptone, 0.5% NaCl and 0.25% agar and the swimming diameter was measured after 6 h at 37 °C. Bars represent mean values with error bars to represent SD from three independent replicates. Differences between mean values were assessed by two-tailed Student’s t-test (ns, not significant; *p < 0.05, **p < 0.01, and ***p < 0.001 compared to *E. coli* MG1655 vector control). VC = pBAD28.

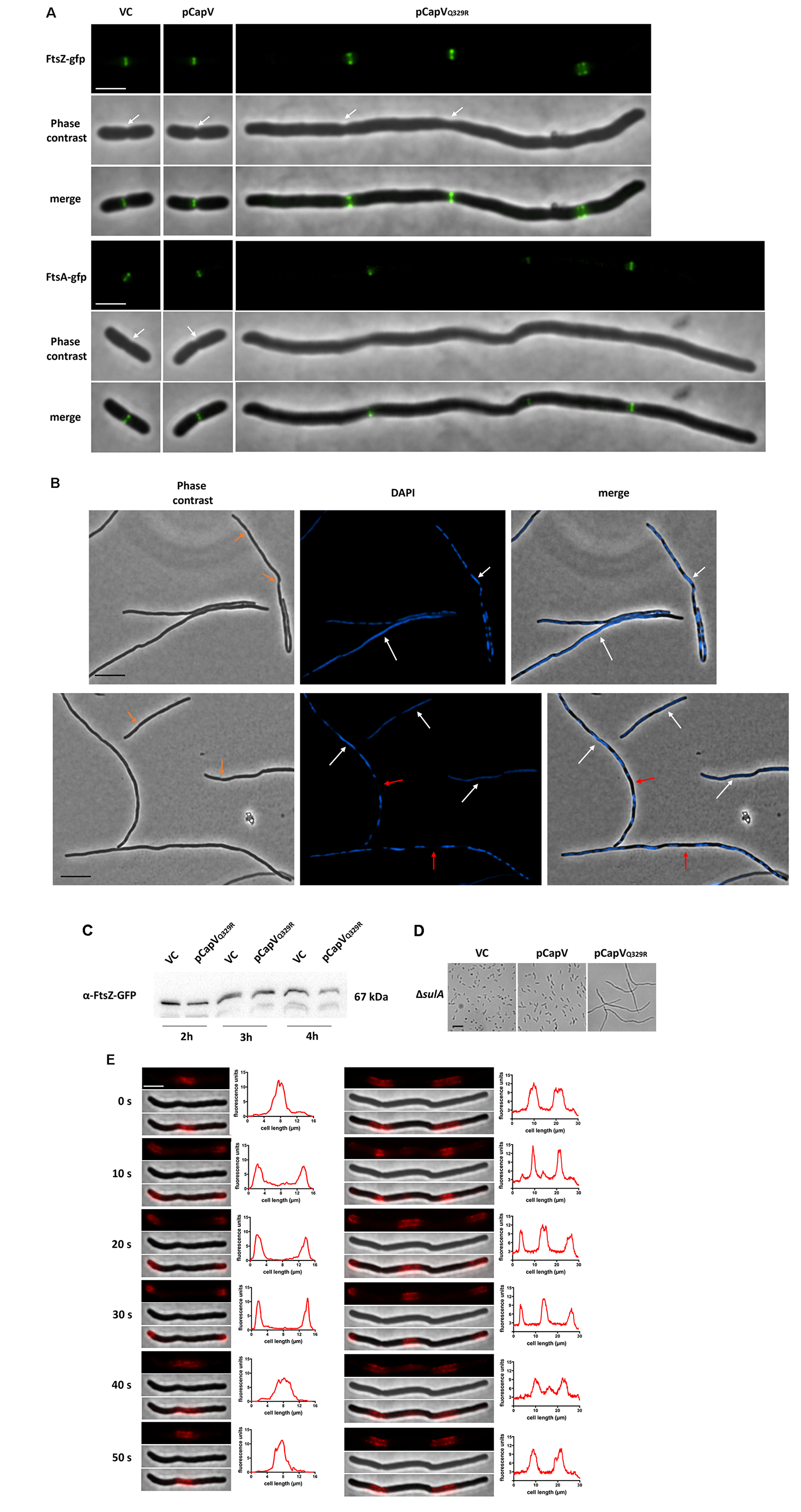

Fig. S4. CapV_Q329R_-induced cell filamentation is independent of *sulA* and does hardly affect FtsZ and FtsA production. (A) Cells were grown for 4 h at 37 °C and placed on an agarose pad to be observed by fluorescence microscopy. Septa are indicated by white arrows. Bar, 3 µm. (B) Cells expressing FtsZ-GFP protein were grown in TB medium at 37 °C, and samples were harvested after 2 h, 3 h, and 4 h of growth for Western blot analysis. (C) Chromosomal segregation is impaired in filamenting cells. Cells were cultured in TB medium at 37 °C for 4 h, stained with DAPI and assessed under fluorescence microscopy immediately. Large fragments of unsegregated nucleoids are indicated by white arrows. Suspected septa are indicated by orange arrows. Large spaces between nucleoids are indicated by red arrows. Bar, 5 µm. (D) CapV_Q329R_-induced cell filamentation is *sulA-*independent. Light microscopy pictures of cell filamentation in an *E. coli* MG1655 Δ*sulA* mutant after 4 h incubation at 37 °C imaged by light microscopy. Bar, 5 µm. (E) Time-lapse analysis of mCherry-MinC expressing cells (PB318) upon CapV_Q329R_ overexpression (see also Fig. 3B). A representative elongating cell is displayed. Graphs on the right of the fluorescence images display the line profiles of fluorescent signals emanating from the cell. Arbitrary fluorescent units are obtained, analyzed by the Fiji ImageJ 1.8.0 software and plotted on the y-axis; cell length in µm is plotted on the x-axis. Bar, 3 µm. VC = pBAD28; CapV = wild type CapV cloned in pBAD28; CapV_Q329R_ = mutant CapV_Q329R_ cloned in pBAD28.

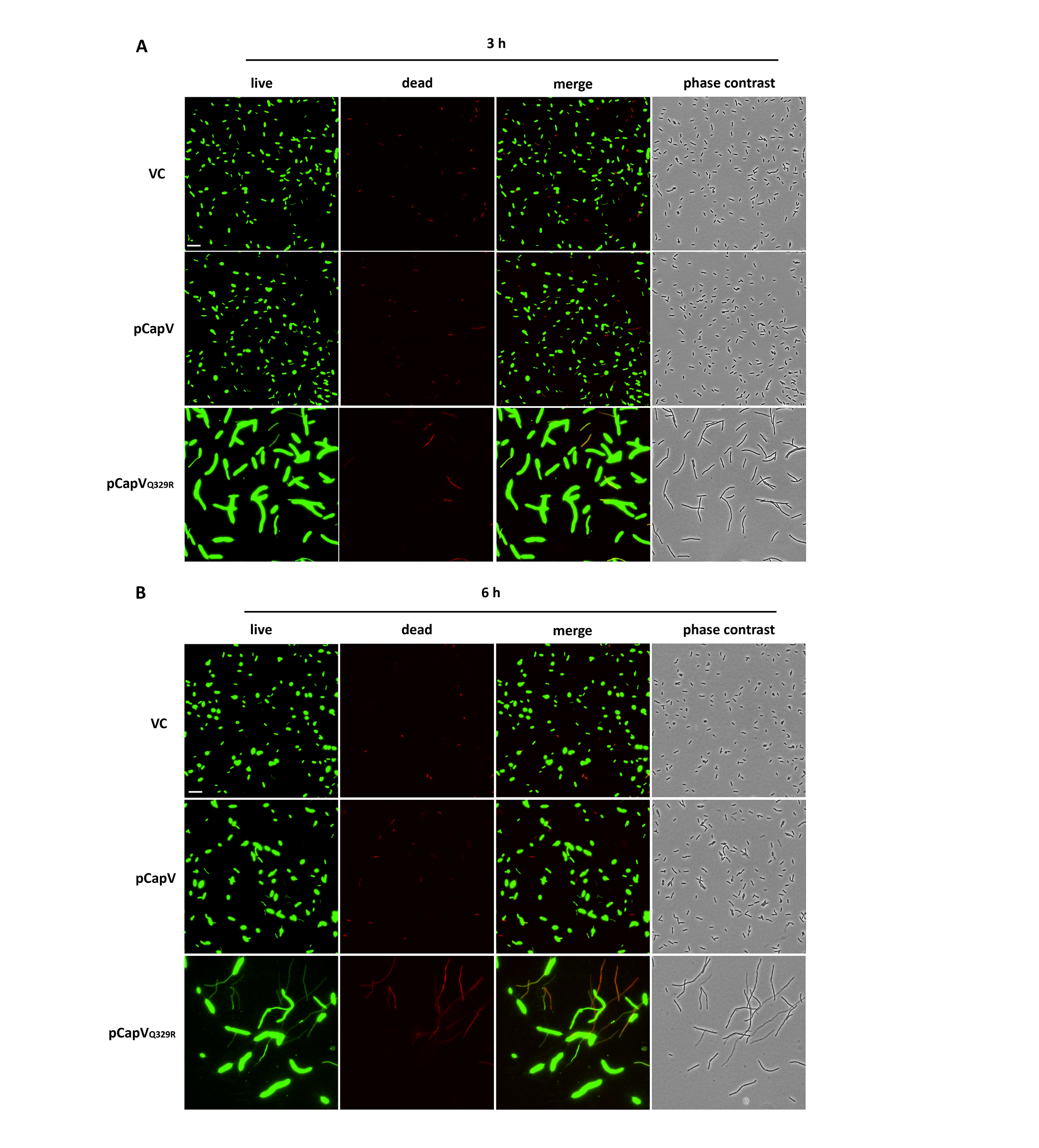

Fig. S5. Cell viability assay upon production of CapV and CapV_Q329R_ by the LIVE/DEAD™ BacLight™ Viability Kit. Cells were incubated for 3 h and 6 h at 37 °C and stained with SYTO 9 and propidium iodide (PI). The cell viability was analyzed under a fluorescence microscope. Green, live cells. Red, dead cells. Bar, 5 µm. VC = pBAD28; CapV=wild type *capV* cloned in pBAD28; CapV_Q329R_ = mutant CapV_Q329R_ cloned in pBAD28.

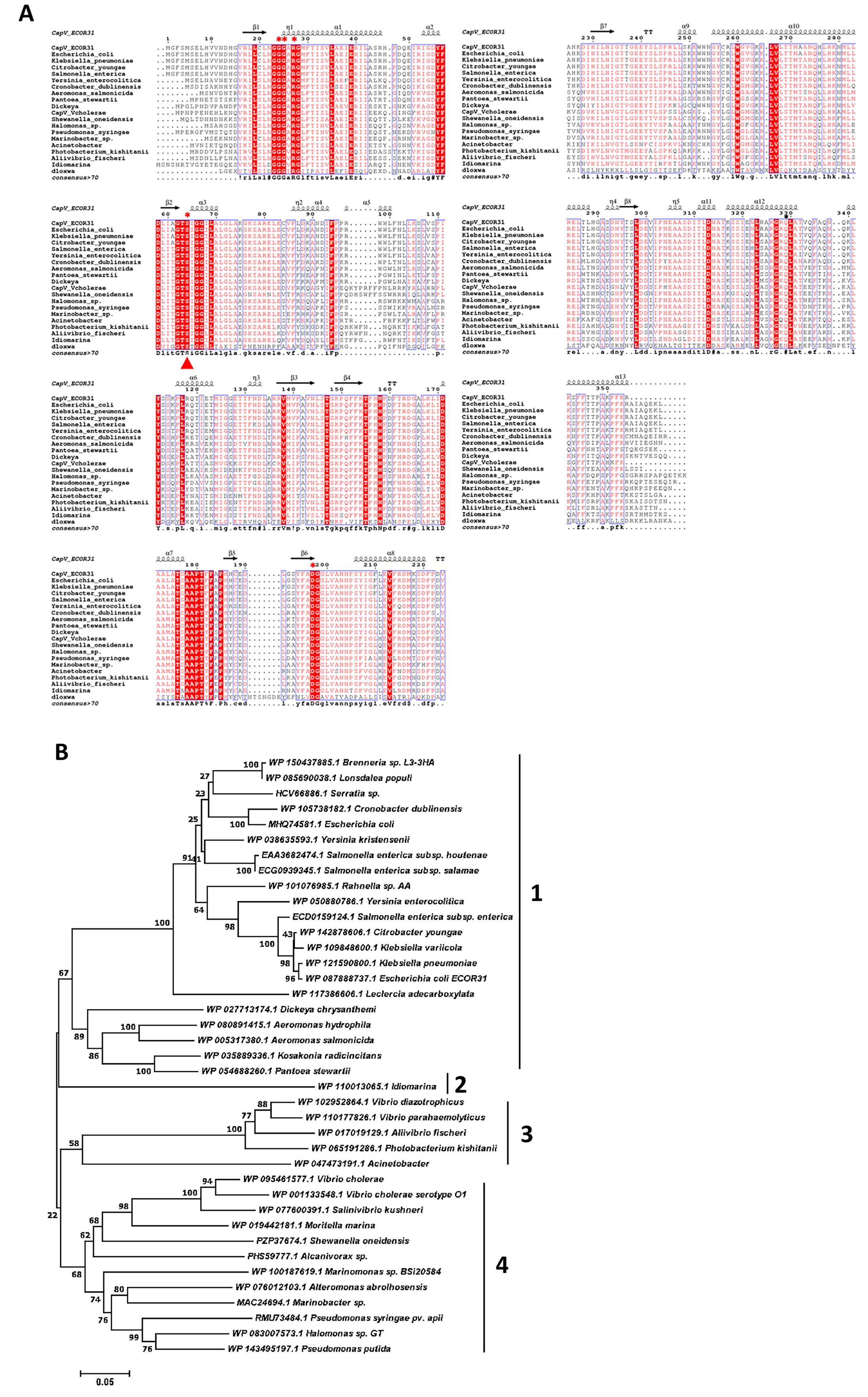

Fig. S6. Phylogenetic and bioinformatic analysis of CapV homologs. (A) Alignment of CapV from *E. coli* ECOR31, *V. cholerae* biovar El Tor and other species with >60% sequence identity. The putative secondary structures of CapV from homology modeling are shown above the alignments. Completely conserved residues are shown in white on a red background. Conserved residues are boxed. The putative catalytic residues of CapV are indicated with filled red triangles. CapV_Q329_ is marked with a black asterisk above the sequence. The residues in CapV_Q329R_ mutated to alanine are marked with red asterisks above the sequence. The consensus sequence below the alignment indicates in uppercase residues with 100% conservation and in lowercase residues higher than >70% conservation. Alignment was performed using CLUSTALW using standard parameters (*4*), and the result processed with ESPript 3.0 (*5*). (B) Maximum likelihood phylogenetic reconstruction of CapV homologs. CapV from *E. coli* ECOR31 was used as the query in the NCBI Blast search and representative proteins (one representative per genus of equal homology) with >60% aa identity over the entire length of the protein sequence were retrieved.

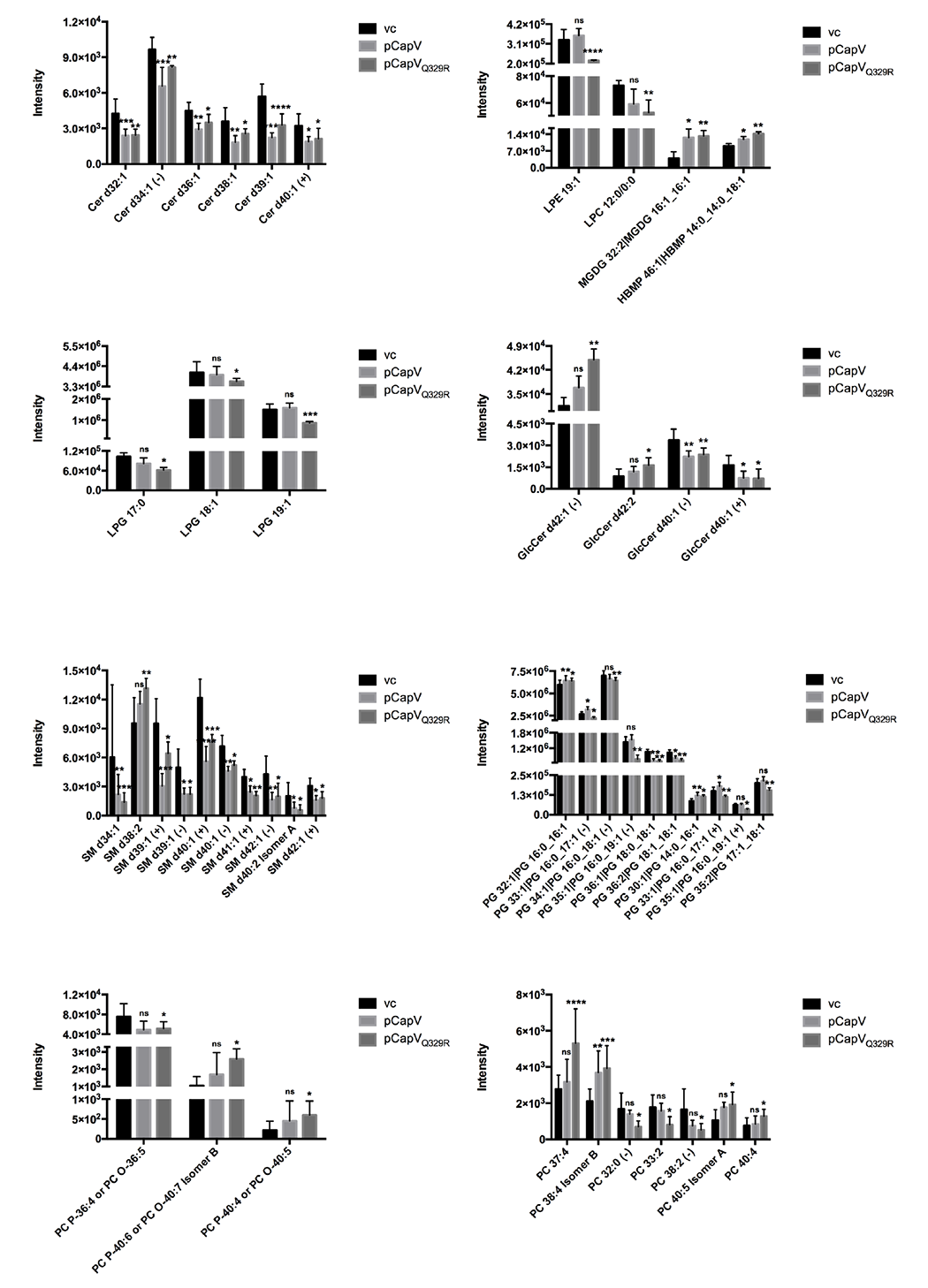

Fig. S7: Lipidomic analysis of *E. coli* MG1655 vector control vc pBAD28 and overexpressing CapV and CapV_Q329R_. Relative abundance of Cer, ceramide; LPG, lysophosphatidylglycerol; SM, sphingomyelin; PC, phosphatidylcholine; LPE, lysophosphatidylethanolamine; LPC, lysophosphatidylcholine; MGDG, monogalactosyldiacylglycerol; HBMP, 1-monoacylglycerol-phospho-2,3-diacylglycerol; GlcCer, glycosylceramide and PG, phosphatidylglycerol derivatives by untargeted CSH-QTOF MS analysis. Bars represent mean values from six independent replicates with error bars to represent SD.

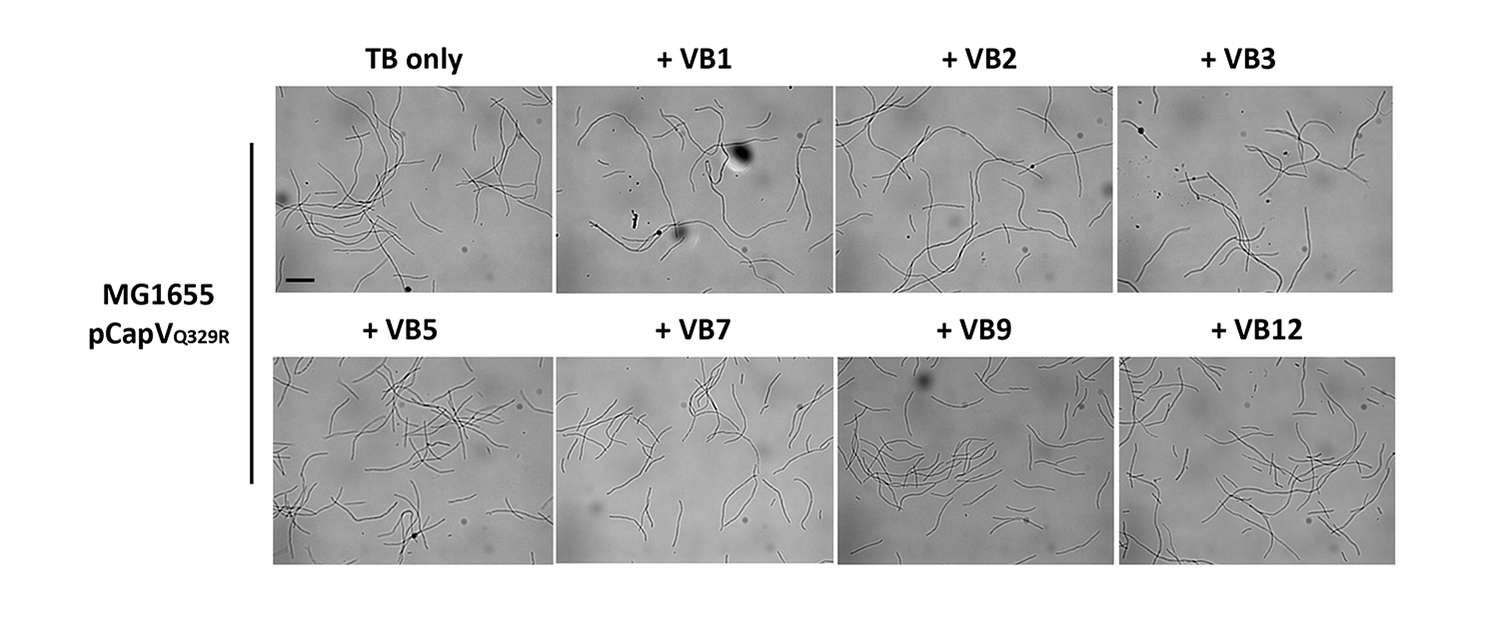

Fig. S8. Effects of various B-vitamins on CapV_Q329R_-induced cell filamentation of *E.coli* MG1655. Addition of Vitamin B1 (thiamine, 0.5%), B2 (riboflavin, 0.5%), B3 (nicotinamide, 0.5%), B5 (pantothenate, 15 mg/ml), B7 (biotin, 100 ug/ml), B9 (folic acid, 80 ug/ml), and B12 (cobalamin, 200 ug/ml) in TB does not affect cell filamentation of *E. coli* MG1655 induced by CapV_Q329R_ after 4 h at 37 °C. Bar, 10 µm. VC = pBAD28; CapV=wild type *capV* cloned in pBAD28; CapV_Q329R_ = mutant CapV_Q329R_ cloned in pBAD28.

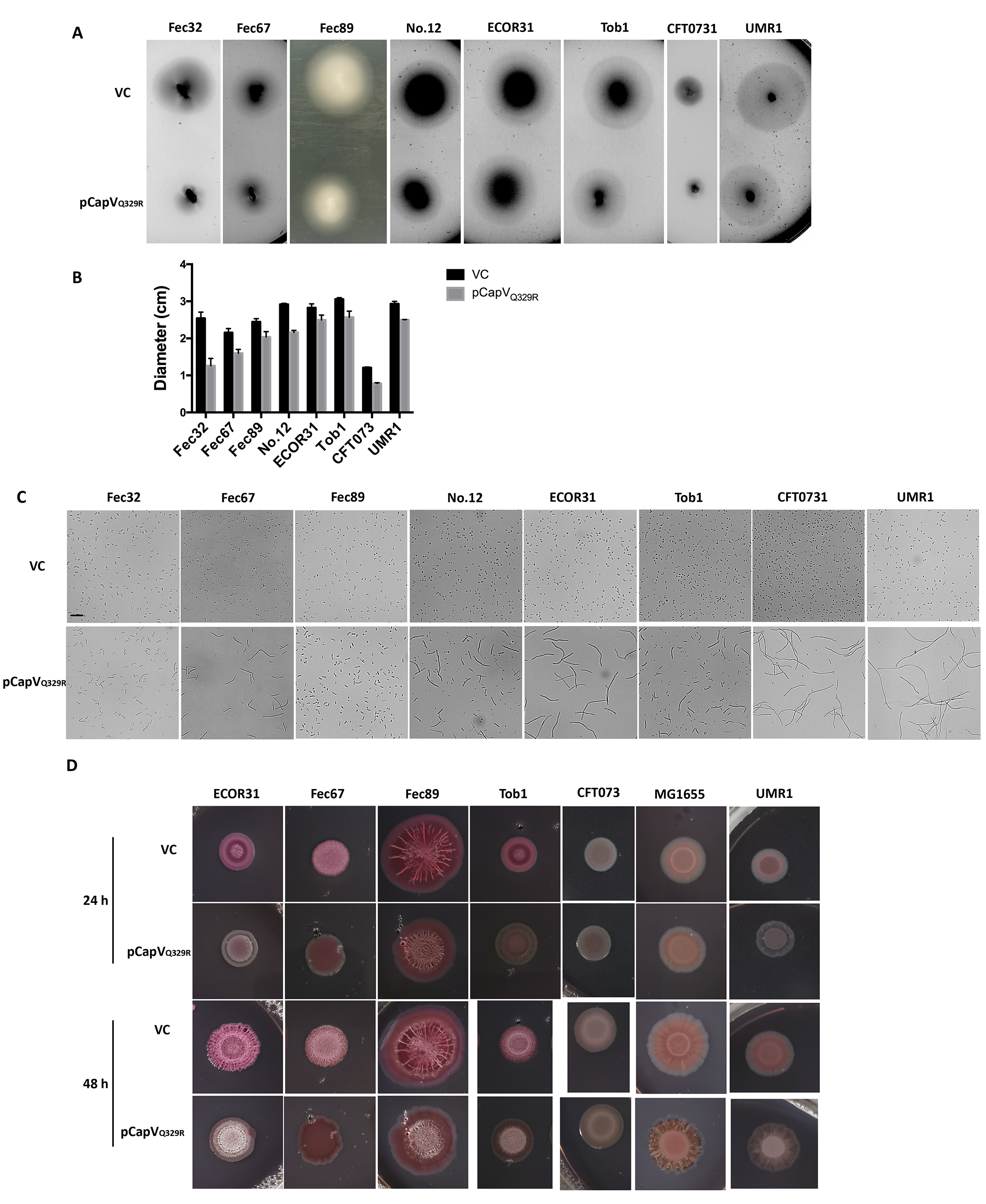

Fig. S9. Apparent swimming motility (A, B), cell filamentation (C ) and rdar biofilm formation (D) of *E. coli* strains Fec32, Fec67, Fec89, No.12, ECOR31, Tob1, CFT073, and *S. typhimurium* UMR1 upon overexpression of CapV_Q329R_. (A-B) 3 µl of OD_600_ = 5 cells were inoculated into soft agar plates containing 1% tryptone, 0.5% NaCl and 0.25% agar. The swimming diameter was measured after 6.5 h for *E. coli* ECOR31 and *S. typhimurium* UMR1, 8 h for *E. coli* Tob1, 9 h for *E. coli* No.12, Fec89, and CFT073, 10 h for *E. coli* Fec67 and 16 h for *E. coli* Fec32, respectively, at 37 °C. Bars represent mean values from three biologically independent replicates. (C ) Cell filamentation of each strain was examined after 6 h of induction at 37 °C by a light microscope. Bar, 10 µm. VC = pBAD28; CapV=wild type capV cloned in pBAD28; CapV_Q329R_ = mutant CapV_Q329R_ cloned in pBAD28. (D) Cells were grown on salt-free LB agar plates at 37 °C and observed after 24 h and 48 h.

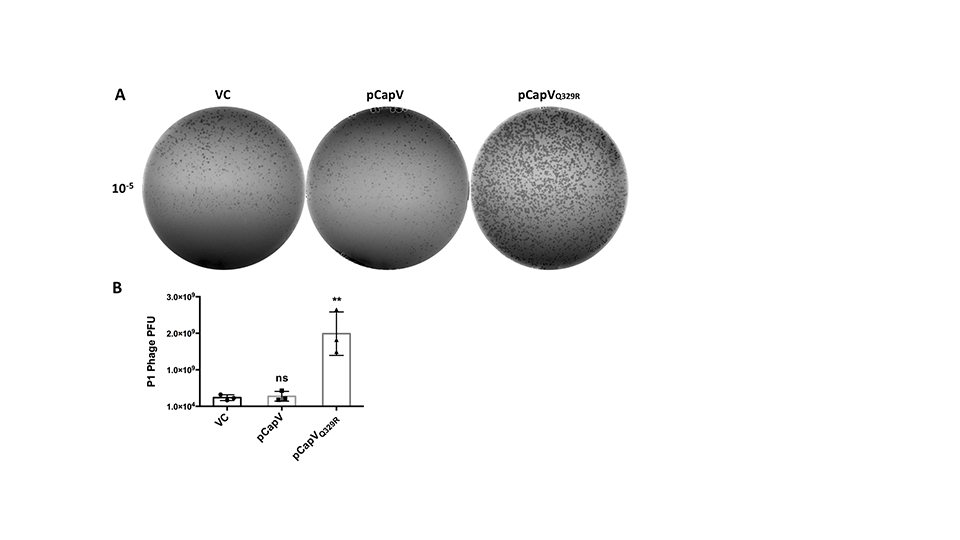

Fig. S10. Plaque formation of bacteriophage P1 on *E. coli* MG1655 vector control and strains overexpressing CapV and CapV_Q329R_. Vector control=pBAD28. (A) Two-layer agar plates with bacteriophage P1 overlay leading to plaque formation on the bacterial lawn. (B) Graph displaying plaque-forming units (PFU) per milliliter; bar graph represents average of three biologically independent replicates.

**Supplementary tables**

**Table S1 Bacterial strains and plasmids used in this study.**

| **Strain or plasmid** | **Genotype or description** | **Reference or source** |
| --- | --- | --- |
| *E. coli* strains | | |
| K-12 derivatives | | |
| MG1655 | ATCC 700926 | *(6)* |
| MG1655 Δ*sulA* | MG1655 Δ*sulA::kanR* | *(7, 8)* |
| BS001 | MG1655 *ftsZ::gfp* | *(9)* |
| PB318 | MG1655 *mCherry-minC (Km^r^) yPet-zapB frt* | *(10)* |
| NEB 5-alpha | High efficiency chemically competent cells, T1 phage resistant and *endA* deficient | New England Biolabs |
| Top10 | F- *mcrA* Δ(*mrr*-*hsdRMS*-*mcrBC*) φ80*lacZ*ΔM15 Δ*lacX74 nupG recA1* *araD139* Δ(*ara-leu*)7697 *galE15 galK16 rpsL*(StrR) *endA1* λ- | Invitrogen |
| Wild type *E. coli* isolates | | |
| ECOR31 | Strain 31 of the *E. coli* reference collection (ECOR) | *(2)* |
| Tob1 | Commensal human faecal isolate | *(11)* |
| Fec32 | Commensal human faecal isolate | *(12)* |
| Fec67 | Commensal human faecal isolate | *(12)* |
| Fec89 | Commensal human faecal isolate | *(12)* |
| No.12 | Wild-type pyelonephritis isolate from urine | Laboratory collection |
| CFT073 | Wild-type pyelonephritis isolate | Laboratory collection |
| *Salmonella enterica* serovar Typhimurium | | |
| UMR1 | ATCC 14028 Nal^r^, rdar at 28 °C | *(13)* |
| *Vibrio cholerae* | | |
| *V. cholerae* C6706 | *V. cholerae* O1 El Tor strain C6706 | Laboratory collection |
| Plasmids | | |
| pBAD28 | Arabinose-regulated promoter; Amp^r^; Cm^r^ | *(14)* |
| p78901 | pBAD28::*capV dncV vc0180 vc0181*; Cm^r^ | *(3)* |
| pDncV | pBAD28::DncV-His_6_; Cm^r^ | *(3)* |
| p78901 | pBAD28::*capV dncV vc0180*; Cm^r^ | This study |
| pCapV | pBAD28::CapV_;_ Cm^r^ | This study |
| pCapV_His | pBAD28:: CapV with C terminal 6X His tag_;_ Cm^r^ | This study |
| pCapV_Q329R_ | pBAD28::CapV_Q329R_; Cm^r^ | This study |
| pCapV_Q329R__His6 | pBAD28::CapV_Q329R_ with C terminal 6X His tag_;_ Cm^r^ | This study |
| pCapV_Q329K_ | pBAD28::CapV_Q329K;_ Cm^r^ | This study |
| pCapV_Q329R/G24A/G25A_ | pBAD28::CapV_Q329R/G24A/G25A;_ Cm^r^ | This study |
| pCapV_Q329R/R27A_ | pBAD28::CapV_Q329R/R27A;_ Cm^r^ | This study |
| pCapV_Q329R/S64A_ | pBAD28::CapV_Q329R/S64A;_ Cm^r^ | This study |
| pCapV_Q329R/D197A_ | pBAD28::CapV_Q329R/D197A;_ Cm^r^ | This study |
| pCapV_Q329R/D197A__His6 | pBAD28::CapV_Q329R/D197A_ with C terminal 6X His tag_;_ Cm^r^ | This study |

**Table S2 Primer used in this study.**

| **Primer** | **Sequence (5’-3’)** |
| --- | --- |
| Gene cloning | |
| VC0178-81-SacI-fw | GGCGAGCTCGCTATATTCTCTGGTTATGGGGTTTTCAATGTCTG^a^ |
| VC0178-XbaI-rv | GCTCTAGATCAGAGTTTCTCCTGCGC^a^ |
| VC0180-XmaI-rv | CCCCCCGGGTCATTTATGTTCATTACACACAGGGC^a^ |
| DncV-SacI-fw | GGCGAGCTCGCAGGAGAAACTCTGATGCCTTG^a^ |
| VC0178-81-SphI-rv | ACATGCATGCTTACTTCTCT CTTATAC^a^ |
| VC0180-SacI-fw | GGCGAGCTCGATGGTAAGTGGCTGATGAAGGAC^a^ |
| DHIII-RV | CCCAAGCTTCCGCATCCCGTCATCACTTGAT^a^ |
| 78_XbaI_fw | GCTCTAGAGCTATATTCTCTGGTTATGGGGTTTTCAATGTCTG^a^ |
| 78-SacI-rv | GGCGAGCTCTCAATGATGATGATGATGATGGAGTTTCTC CTGCGC^a^ |
| Site-directed mutagenesis | |
| CapV-Q329K-fw | AAGGGGAAGAaAACTCGCTAC |
| CapV-Q329K-rv | GCGCGCAGGTTTTCAATG |
| G24/25A-fw | CTAAATGGTGcggcGGCCAGAGGGATGTTTAC |
| G24/25A-rv | GCAGAGTAGCCGTACTCC |
| R27A-fw | TGGTGGGGCCgcgGGGATGTTTAC |
| R27A-rv | CCATTTAGGCAGAGTAGC |
| CapV-S64A-Q5-fw | CGCAGGGACAgCGATAGGTGG^b^ |
| CapV-S64A-Q5-rv | ATGAGGTCAAAGTAATCGCCAATTC |
| CapV-D197A-Q5-fw | TATTTTGCTGcgGGAGGTCTGGTC^b^ |
| CapV-D197A-Q5-rv | AGAACCAAGGTCTTCACAATG |
| d-xbaI-fw | ATGGTGAGCAAGGGCGAG |
| d-xbaI-rv | GAGTTTCTCCTGCGCGATTG |
| DncV-D129/131A-Q5-fw | tgcgGATGGCACCTACATGCCC^b^ |
| DncV-D129/131A-Q5-rv | atcgCCATTTCCTGCCCCGGAT^b^ |
| DncV-D197A-Q5-fw | ACACACATTGcgGTACCGATGTATG^b^ |
| DncV-D197A-Q5-rv | TTTCTCCCGATAGATTTTG |
| VC0180-C91A-Q5-fw | AGGGTTCCTTgcgTATGTTGAGCAGATGGAAGCAGACTGGG^b^ |
| VC0180-C91A-Q5-rv | TCCAGGGCGACATGGGGC |
| VC0180-C355A-Q5-fw | GTTGATAGGTgcgGGAACAATTGGAGGG^b^ |
| VC0180-C355A-Q5-rv | GCTATACGCTTTGTTGAAAG |
| VC0181-E123A-Q5-fw | TATTTAGGCGcgTGGCATACACATC^b^ |
| VC0181-E123A-Q5-rv | AACAAGAAAGCCATTTGAC |
| 2401-fw | AGGGGAAGACaACTCGCTACTG^b^ |
| 2401-rv | TGCGCGCAGGTTTTCAAT |
| 2805-fw | GACTTCATCAaATTCCAGCCTC^b^ |
| 2805-rv | ATCGCGTGCTTCATCTTC |
| 3324-fw | GGCGGGCATCtGCGTTCAGTT^b^ |
| 3324-rv | GATACGTTTACAGCTTTCGTTGAACC |
| 5496-fw | ATATCCTTGAgGGGGGGCGTT^b^ |
| 5496-rv | GTTCATTGTAAGTCCACTTTAACTC |
| Confirmatory primers | |
| pBAD30-fw | GTCTATAATCACGGCAGAAAAGTCCAC |
| pBAD30-rv | CTGTTTTATCAGACCGCTTCTGC |
| VC0178-inside-fw | GAAACGATGATCGGTGGTGAG |
| DncV-seq | TACATGCCCATGACGGTGTT |
| VC0180-inside-fw | CCCATCAGCACGCATTACATTTC |
| 80-inside-fw2 | GATACACCAGACCCTCATTG |
| VC0181-inside-rv | GGATGTGTATGCCATTCGCCTA |

Underlined indicates ^a^restriction sites, ^b^mutated codons.

**Supplementary Figures**

Additional file types that cannot be embedded into the Word file:

Movie S1 Movie of *E. coli* MG1655 cells upon overexpression of CapV wild type and CapV_Q329R_ induced by 0.1% L-arabinose at 37 °C in TB medium while being examined after 4 and 6 h under a microscope. VC = pBAD28; CapV=wild type capV cloned in pBAD28; CapV_Q329R_ = mutant CapV_Q329R_ cloned in pBAD28.

Movie S2. *E. coli* MG1655 cells examined under a light microscope 22 h after induction of CapV_Q329R_ production by 0.2% L-arabinose in TB medium at 37 °C.
